# Long-term adaptation to galactose as a sole carbon source selects for mutations in nutrient signaling pathways

**DOI:** 10.1101/2022.05.17.492354

**Authors:** Artemiza A. Martínez, Andrew Conboy, Sean W. Buskirk, Daniel A. Marad, Gregory I. Lang

## Abstract

Galactose is a secondary fermentable sugar that requires specific regulatory and structural genes for its assimilation, which are under catabolite repression by glucose. When glucose is absent, the catabolic repression is attenuated, and the structural *GAL* genes are fully activated. In *Saccharomyces cerevisiae*, the *GAL* pathway is under selection in environments where galactose is present. However, it is unclear the adaptive strategies in response to long-term propagation in galactose as a sole carbon source in laboratory evolution experiments. Here, we performed a 4,000-generation evolution experiment using 48 diploid *Saccharomyces cerevisiae* populations to study adaptation in galactose. We show that fitness gains were greater in the galactose-evolved population than in identically evolved populations with glucose as a sole carbon source. Whole-genome sequencing of 96 evolved clones revealed recurrent *de novo* single nucleotide mutations in candidate targets of selection, copy number variations, and ploidy changes. We find that most mutations that improve fitness in galactose lie outside of the canonical *GAL* pathway and are involved in nutrient signaling. Reconstruction of specific evolved alleles in candidate target of selection, *SEC23* and *IRA1*, showed a significant increase in fitness in galactose compared to glucose. In addition, most of our evolved populations (28/46; 61%) fixed aneuploidies on Chromosome VIII, suggesting a parallel adaptive amplification. Finally, we show greater loss of extrachromosomal elements in our glucose-evolved lineages compared with previous glucose evolution. Broadly, these data further our understanding of the evolutionary pressures that drive adaptation to less-preferred carbon sources.

## INTRODUCTION

Catabolic repression–the inability to utilize a less-preferred carbon source when a more favorable one is present–is a core principle of metabolic gene regulation. As was first demonstrated for the *lac* operon in *E. coli*, expression of the structural genes needed to assimilate alternative carbon sources require two conditions be met: the absence of the preferred carbon source and the presence of the alternative carbon source (Jacob and Monod 1961). This logic is used to regulate the assimilation of a variety of secondary fermentable (e.g., sucrose, maltose, and galactose) and non-fermentable (e.g., acetate, glycerol, and ethanol) carbon sources (Zaman et al. 2008; Fendt and Sauer 2010).

In the yeast, *Saccharomyces cerevisiae*, galactose is a widely used secondary carbon source. Galactose is a common sugar found in dairy products, fruits, grains, and vegetables (Acosta and Gross 1995; Lee et al. 2011). Galactose assimilation requires three regulatory genes and four structural genes, which are expressed only when glucose is absent and galactose is present. In the absence of glucose, the Snf1 complex attenuates catabolite repression by phosphorylating the Mig1 transcriptional repressor, leading to its inactivation (Conrad et al. 2014; Nair and Sarma 2021). Full activation of galactose structural genes is mediated by a transcription factor (Gal4p), a repressor (Gal80p), and a co-inducer (Gal3p). In the absence of galactose, Gal80p sequesters Gal4p in the cytoplasm. When present, galactose binds to Gal3p, which in turn binds to Gal80p, releasing Gal4p. Once released, Gal4p relocalizes to the nucleus and increases up to ∼1000-fold the transcription of the *GAL* structural genes, which are clustered in a 7 kb region on Chromosome II (Sellick et al. 2008; Conrad et al. 2014).

Galactose initially enters the cell through low-affinity hexose transporters (Escalante-Chong et al. 2015). However, once the switch occurs, galactose enters the cell predominantly through the high-affinity galactose transporter Gal2 (Conrad et al. 2014). Once in the cell, three main enzymes are necessary to catalyze four sequential steps in galactose assimilation. Gal1p is the galactokinase. Gal10p contains two catalytic domains: a mutarotase that interconverts galactose enantiomers and an epimerase that converts UDP-galactose to UDP-glucose. Gal7p is the galactose-1-phosphate uridylyltransferase. These enzyme are tightly regulated to reduce their costly expression and avoid accumulation of the toxic intermediate galactose-1-phosphate (Slepak et al. 2005; Wang et al. 2015).

The genetic switch from glucose to galactose utilization has been extensively studied as a model system to understand evolution of gene clustering (Hittinger and Carroll 2007; Lang and Botstein 2011; Harrison et al. 2021), transcriptional regulation (Brink et al. 2009; New et al. 2014; Peng et al. 2015), nutrient-sensing metabolism (Escalante-Chong et al. 2015; Lee et al. 2017), and cellular memory (Acar et al. 2005; Sood and Brickner 2017). The clustering and regulation of the *GAL* pathway has evolved independently multiple times in fungi. Relative to other yeast, *S. cerevisiae* evolved strong repression and slow induction of the *GAL* genes, which confers a fitness advantage in environments where glucose is in excess (Harrison et al. 2021). However, studies on natural environments identified novel strategies in *S. cerevisiae* isolates that are associated with improved growth on galactose. For example, natural variation in catabolite repression varies with the length of diauxic lag such that galactose is only assimilated once glucose drops below a certain threshold (Wang et al. 2015). Lineages isolated from dairy products harbor *GAL* variation, such as, *GAL3* polymorphisms, *GAL2* amplification, and *GAL* cluster introgressions from *S. paradoxus* reflecting adaptation to a galactose-rich environment (Duan et al. 2018; Legras et al. 2018; Boocock et al. 2021). These suggest that the structural and regulatory genes in the *GAL* pathway are under selection in environments where galactose is present. However, natural environments are complex and variable environments, therefore it is unclear how selection acts to improve growth in a simple environment containing galactose as the sole-carbon source.

Experimental evolution of microbes is a powerful tool to identify genomic changes during adaptation in different environments (Voordeckers et al. 2015; Swamy and Zhou 2019). The molecular targets of selection can be informative on the pressure that drive adaptation in each system. For example, in glucose-limitation laboratory evolution, recurrent mutations arise in nutrient signaling pathways, such as the Ras/PKA pathway and TOR pathway. In chemostat, amplification of glucose transporters is another recurrent adaptive mechanism (Kao and Sherlock 2008; Wenger et al. 2011; Venkataram et al. 2016). In glucose-rich laboratory evolution, genetic targets of selection are involved in cell wall biogenesis, assembly, and cytokinesis, as well as nutrient sensing and signaling (Lang et al. 2013; Fisher et al. 2018; Marad et al. 2018; Johnson et al. 2021). Previous galactose, laboratory-evolution studies identified a mutation in transcriptional repressor *GAL80* (Quarterman et al. 2016), in proteins involved in polarized growth (Jerison et al. 2020), and in the Ras/PKA pathway (Hong et al. 2011a; Hong and Nielsen 2013). In contrast to variation found in galactose-rich natural environments, mutations identified in laboratory evolution are predominantly outside the canonical *GAL* genes.

Previously, we evolved 48 diploid populations in glucose over 4,000 generations (Marad et al. 2018). Here, we perform an identical 4,000 generation diploid evolution experiment using galactose as the solely carbon source. We directly compare the rates of adaptation and genetic targets of selection between glucose and galactose evolution. Consistent with previous work, we find that most mutations that improve fitness in galactose lie outside of the canonical *GAL* pathway and are involved in nutrient signaling. In addition, we identify recurrent copy number variants, including aneuploidies on Chromosome VIII. Reconstruction of specific evolved alleles in *SEC23* and *IRA1* showed significant increases in fitness under galactose conditions. Finally, we show multiple losses of extrachromosomal elements in most of our evolved lineages compared with glucose evolution. These data can be explored further to better understand the evolutionary pressure that could drive adaptation under less-preferred carbon source in long-term experimental evolution.

## RESULTS

### Higher rate of adaptation in rich galactose medium compared to rich glucose medium

Previously, we evolved 48 diploid populations for 4,000 generations in rich glucose (YPD) medium (Marad et al. 2018). To determine the evolutionary response to constant growth in an alternative carbon source, we evolved 48 replicate populations of the same diploid ancestor in a rich medium with galactose as the sole carbon source (YPGal). After 2,000 generations, all populations increased in fitness compared to the ancestor (Figure 1A) with a mean fitness gain of 8.2 ± 0.1% (α= 0.05). This is significantly greater than the fitness gains for the identically evolved populations using glucose as a sole carbon source (*p* < 0.0001, Wilcoxon Rank-Sum test; Figure 1B), which showed mean fitness gains of 3.7 ± 0.06% (α= 0.05) in the first 2,000 generations of evolution.

**Figure 1.**
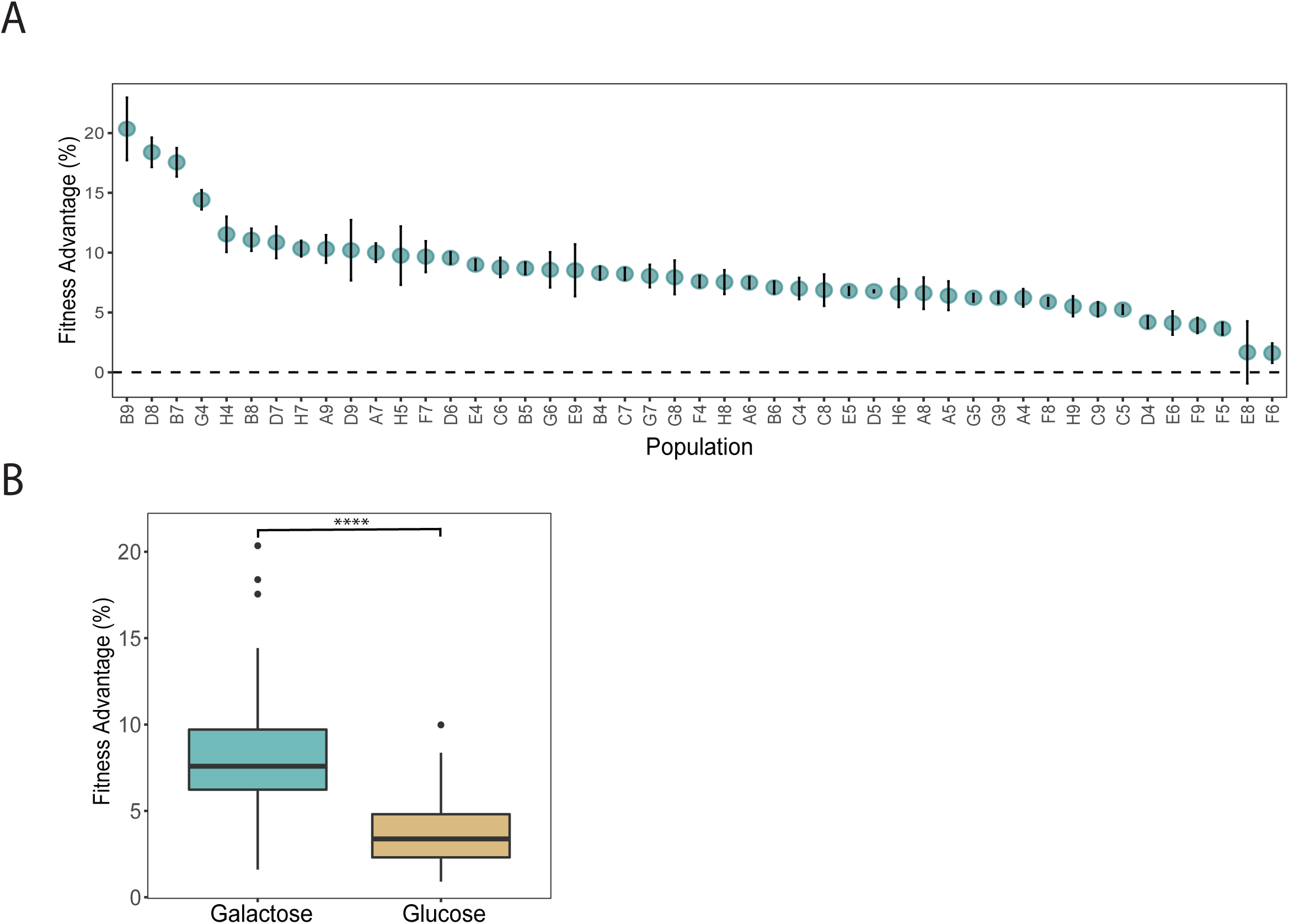
Galactose-evolved populations have a higher fitness advantage after 2,000 generations of laboratory evolution. (A) Relative fitness advantage of 48 evolved populations on galactose. Vertical error bars reflect ± standar error of the regression. (B) Adaptation in galactose has a significantly higher fitness advantage compared with glucose-evolved populations (*p* < 0.0001, Wilcoxon rank-sum test). Box plot reflects the mean fitness of 48 populations in both conditions. Galactose populations have a mean fitness advantage of 8.2%, whereas glucose populations have a mean of 3.7%. Glucose fitness data is from Marad et al, 2018.

### Beneficial mutations are commonly found outside the canonical *GAL* pathway

To identify mutations that arose in the galactose-evolved populations, we sequenced two clones from each of the 48 populations at Generation 4,000. We found 1,529 mutations distributed across all 16 chromosomes with a mean of 33 ± 9 mutations per clone. Of these mutations, 793 are present in both clones from the same population, and 736 mutations are only found in one clone. Of the 1,529 mutations, 91% are single nucleotide polymorphisms, most of which are in coding genes (1092 out of 1529; 71.4%). 712 of these mutations are mostly missense (46.6%), 225 are synonymous (14.7%), 76 are nonsense (5%), 59 are frameshift (3.9%), and 20 are complex mutations (1.3%) (Figure 2 and Supplementary Data). Consistent with our previous work (Fisher et al. 2018; Marad et al. 2018), we find that heterozygous mutations outnumber homozygous mutations ∼19:1 (Supplementary Figure 1A), and that homozygous mutations are clustered on the right arm of Chromosome XII (Supplementary Figure 1B). This region is a hotspot for loss of heterozygosity (LOH) due to the rDNA locus: approximately one third of the homozygous mutations are located on this chromosome.

**Figure 2.**
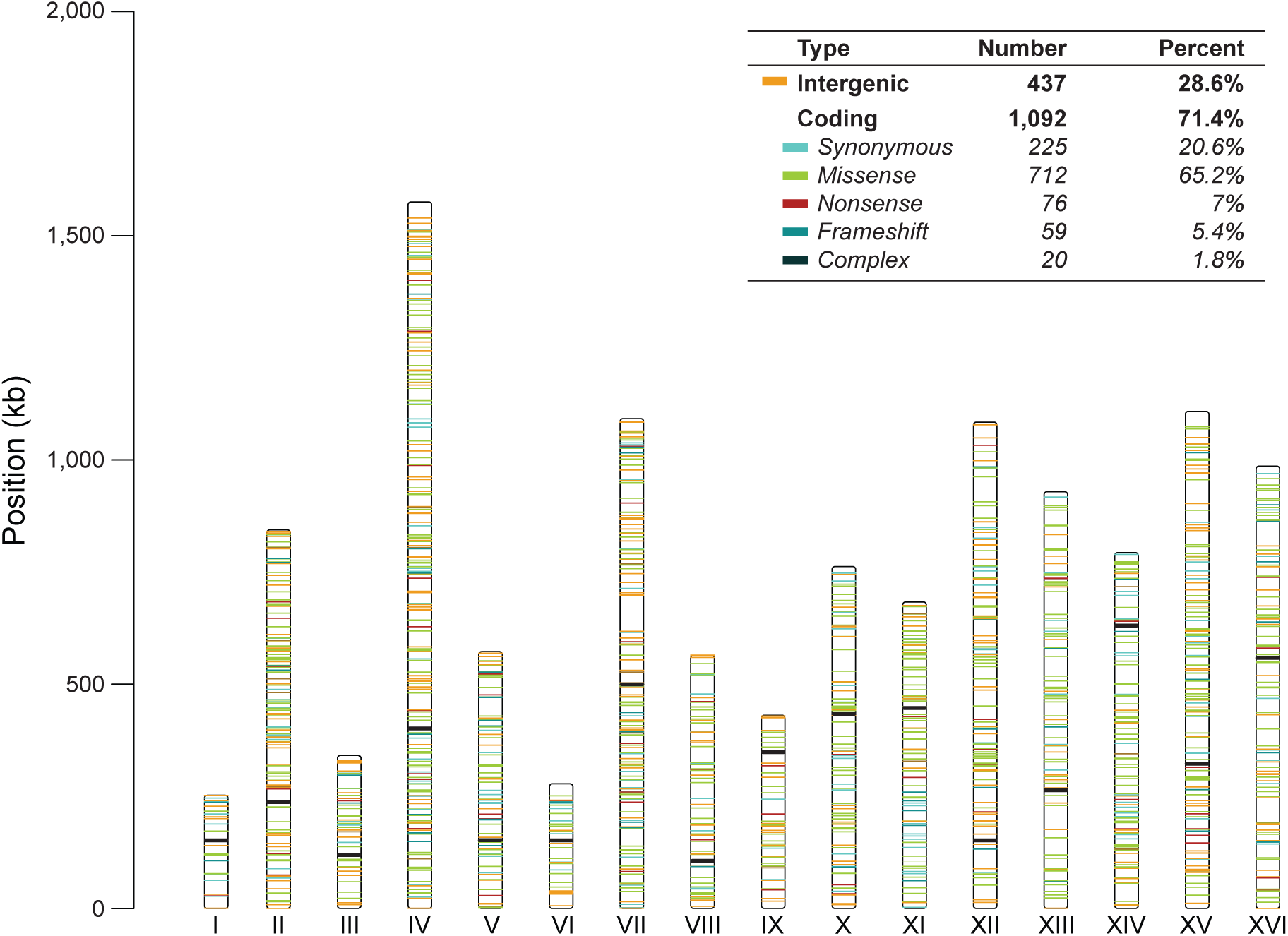
The spectrum of mutations in 96-evolved clones under galactose evolution. Distribution of 1,529 evolved mutations across 16 Chromosomes. The vertical bars represent each Chromosome labeled by Roman numeral. The horizontal lines reflect each mutation colored based upon their protein-coding effect (top box). Other mutations group conservative inframes, disruptive inframes, and complex mutations. Centromers are represented with a black horizontal line.

Genes that fixed mutations more often than expected by chance are candidate targets of adaptation. We identified 13 targets of selection with a *p* < 0.005 including five with *p* < 0.001 (Poisson distribution) (Figure 3A). These genes are principally involved in the Ras/PKA pathway, histone deacetylation, endosome organization/vacuolar biogenesis, and secretory pathway. Notably, we did not find a significant number of mutations in genes with an obvious connection to galactose metabolism. We found three mutations in *GAL10* (two missense and one conservative inframe), two mutations in *GAL1* (one nonsense and one synonymous), two missense mutations in *GAL2*, one missense mutation in *GAL4*, and one missense mutation in *GAL7*. However, none of these mutations were identified as candidates targets of selection at *p* < 0.001.

**Figure 3.**
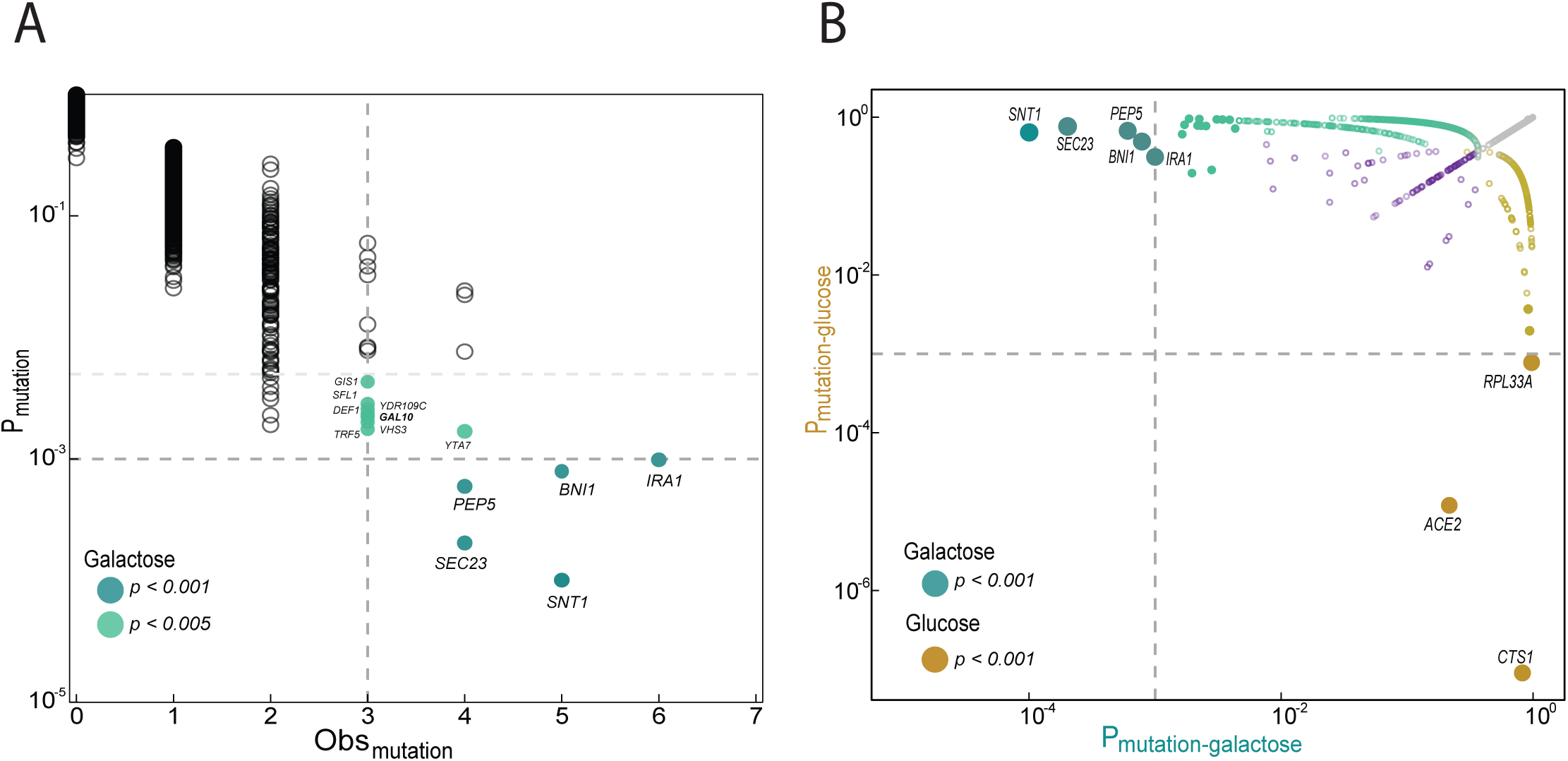
Common targets of selection in galactose-evolved populations. (A) Plotted on the x-axis is the observed number of coding sequence (CDS) mutations in each of the 5800 genes. The y-axis is the probability that the observed number of CDS mutations in each gene occurred by chance. We used a Poisson distribution weighted for the length of each gene. (Fisher et al. 2018). Common targets of selection are genes with 3 or more CDS mutations with corresponding *p*-value< 0.005 (solid light green) and *p*-value< 0.001 (solid dark cyan). (B) Common targets of selection in galactose-evolved populations differ from glucose-evolved populations. Plotted is the ratio of the probability that the observed number of CDS mutations in a gene occurred by chance in galactose populations (cyan circles) versus glucose populations (gold circles). Solid circles indicate the *p*-value< 0.005. Genes with a *p*-value< 0.001were labeled with the gene name and bigger circles. Open purple circles indicate CDS mutated in both conditions. Grey circles indicate CDS that were not mutated under any conditions. Glucose data is from Marad et al. 2018.

To determine if adaptation to galactose occurs through similar genes and pathways as adaptation to glucose, we compared the candidate targets of selection in galactose with our previous data from glucose-evolved populations (Marad et al. 2018). Indeed, none of the candidate targets of selection in galactose are significantly overrepresented in glucose evolution experiment and vice versa (Figure 3B). The most common targets of selection in glucose-evolved populations are cell wall-associated genes, *CTS1, ACE2*, and *RPL33A (p* < 0.001, Poisson distribution; Figure 3B). Of these, only one mutation in *ACE2* was observed in one galactose-evolved population.

Based on the types of mutations observed, we can determine how selection is acting on the candidate targets of selection (Table 1). We observed enrichment for nonsense and frameshift mutations (high impact mutations) in *SEC23, PEP5*, and *IRA1*, suggesting selection for loss-of-function. In contrast, we only observe missense mutations in *SNT1*. Of the five mutations found in *BNI1*, three are synonymous and conservative in-frame mutations, suggesting that this locus may not have a strong effect in the adaptation of galactose. The majority of mutations located in our targets are heterozygous except for *IRA1*, where 2 out of 6 mutations are homozygous.

**Table 1.** Common targets of selection in galactose populations

| Gene | Amino acid change | Zygosity | Clones | Biological process (SGD) |
| --- | --- | --- | --- | --- |
| *SNT1*<br>(Chr III) | Ala168Ser<sup>†</sup> | Heterozygous | **F51, F52** | Subunit of the Set3C deacetylase complex |
|  | Val237Ile | Heterozygous | B42 |  |
|  | Val374Leu | Heterozygous | **E81, E82** |  |
|  | Arg1009Lys<sup>†</sup> | Heterozygous | **A91, A92, E52** |  |
| *SEC23*<br>(Chr XVI) | Gln167* | Heterozygous | **F41, F42** | COPII vesicle coating |
|  | Gln494* | Heterozygous | **A91, A92** |  |
|  | Arg592Ile<sup>†</sup> | Heterozygous | **G61, G62** |  |
|  | Leu701fs | Heterozygous | **H61, H62** |  |
| *PEP5*<br>(Chr XIII) | Tyr406Tyr | Heterozygous | **G61, G62** | Endosome organization<br>/vacuolar organization |
|  | Lys532* | Heterozygous | H42 |  |
|  | Glu650* | Heterozygous | B82 |  |
|  | Tyr808*<sup>†</sup> | Heterozygous | **H51, H51** |  |
| *IRA1*<br>(Chr II) | Ser700Pro<sup>†</sup> | Homozygous | **C41, C42** | Inhibitory Regulator of the RAS-cAMP pathway |
|  | Gln933* | Homozygous | **H81, H82** |  |
|  | Pro1013fs | Heterozygous | **A91, A92** |  |
|  | Leu1402* | Heterozygous | **G91, G92** |  |
|  | Gln2083* | Heterozygous | **A61, A62** |  |
|  | Ala2292Ser | Heterozygous | **B51, B52** |  |

Bolded clones indicate the SNP was found in all clones of the population
\*Nonsense
fs Frameshift
<sup>†</sup> Reconstructed mutations

### Galactose-evolved alleles show carbon source dependence effect

To quantify the fitness effect of individual mutations, we reconstructed candidate evolved mutations in the ancestral background. We selected mutations that are nonsynonymous and present in both clones from their respective populations; *snt1-*A168S, *snt-*R1009K, *sec23*-R592I, *ira1-*S700P (Table 1). As a control, we used two previously identified adaptive mutations, *hsl1*-A262P (Buskirk et al. 2017; Vignogna et al. 2021), and *ace2-*R669* (Marad et al. 2018) from haploid-glucose evolution and diploid-glucose evolution, respectively. Each mutation was reconstructed as haploids, heterozygous diploid, and homozygous diploids. We performed competitive fitness assays of the reconstructed strains against a fluorescently labeled ancestral reference in both galactose and glucose. For all evolved mutations, we find that the fitness effects in glucose and galactose are highly correlated (*r* = 0.69, *p* = 0.0122, Pearson correlation coefficient; Supplementary Figure 2). Nevertheless, the magnitude of the fitness effect was greater in the condition in which the specific mutation arose, and this difference is more exaggerated for homozygous mutations. We found a fitness advantage of the heterozygous *sec23-*R592I mutation of 3.9 ± 0.4% (α= 0.05; *p* < 10^−6^ one-way ANOVA, Tukey’s HSD test) in galactose than glucose and a larger fitness advantage of 6.2 ± 0.5% (α= 0.05; *p* < 10^−8^, one-way ANOVA, Tukey’s HSD test) as homozygous in galactose compared to glucose. We observed similar fitness advantage for *ira1-*S700P mutation; 3.3 ± 0.3 % (α= 0.05) as heterozygous and 5.3 ± 0.3% (α= 0.05) as homozygous (Figure 4B). Despite the greater fitness benefit of the homozygote mutations, only *ira1-*S700P lost heterozygosity in the evolution experiment. Surprisingly, neither of the two *SNT1* alleles provided a selective benefit in galactose or glucose. The heterozygous *snt1-*R1009K mutation is strongly deleterious, but it is neutral as a homozygote. The *snt1-*A592S shows the opposite pattern: the heterozygous mutation is neutral but homozygous mutation is deleterious (Figure 4). As expected, the ploidy, glucose-evolved alleles of *hsl1*-A262P and *ace2-*R669* are more beneficial in glucose although both alleles also improved fitness in galactose (Figure 4 and Supplementary Figure 3). Our results show that evolved mutations on *SNT1* and *IRA1* are beneficial only in the condition in which they arose.

**Figure 4.**
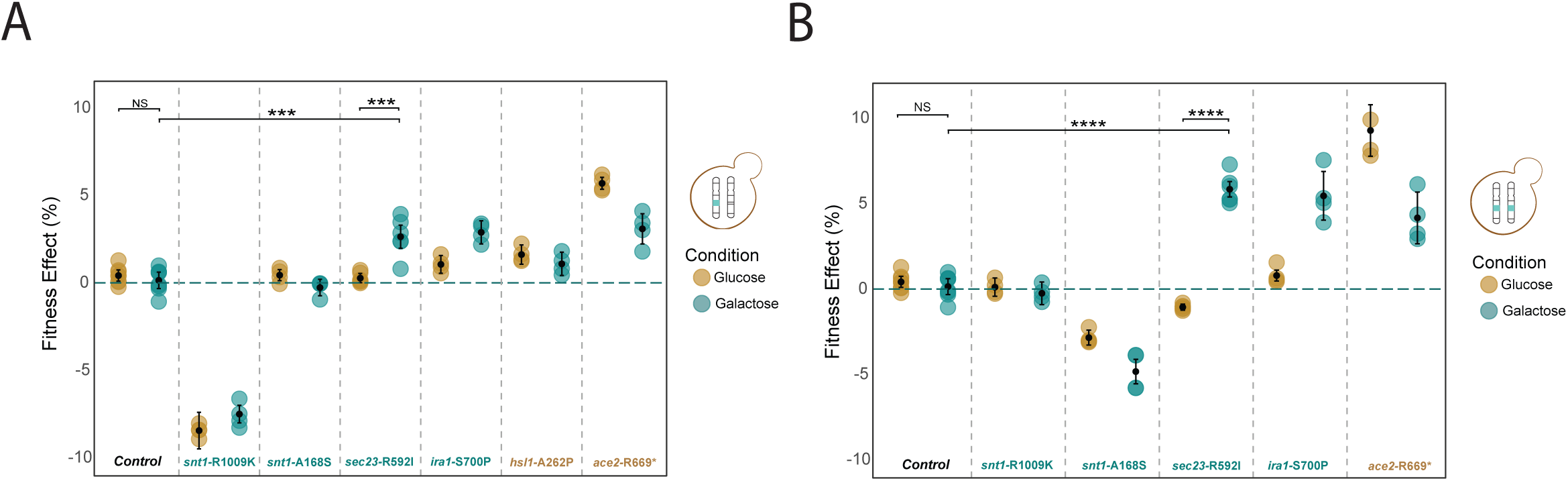
Galactose-evolved mutations show variation in fitness effect and carbon source dependence. Average fitness effects of the reconstructed mutations in *snt1* (*n*=4), *sec23* (*n*=6), and *ira1* (*n*=4) in both zygosities; (A) heterozygous and (B) homozygous mutations. Fitness effect were compared with wild-type version (*n*=6), *hsl1* mutation (*n*=4) from glucose, haploid-evolution (Fisher et al, 2018), and *ace2* mutation (n=4) from glucose, diploid-evolution (Marad et al, 2018). We were not able to generate *hsl1* homozygous mutation. The *sec23*-R592I has a significant fitness gain in galactose compare to glucose. Asterisk (****) indicate *p* < 10^−8^, (***) *p* < 10^−6^, and NS: not significant; one-way ANOVA, tukey’s HSD test. *ira1*-A168S follows the same effect as *sec23-*R592I. We estimated *p*-values only for genes with 6 biological replicates. Error bars are the s.e.m.

### Aneuploidies and CNVs are common in galactose-evolved populations

In addition to point mutations, copy number variants (CNVs) and aneuploidies can contribute to adaptation (Yona et al. 2012; Sunshine et al. 2015; Fisher et al. 2018; Marad et al. 2018). To identify CNVs and aneuploidies in our galactose-evolved populations, we compared the coverage across 16 chromosomes. We find that most of our evolved clones (28/46; 61%) have one or two extra copies of Chromosome VIII. In addition, we find one population with monosomy for Chromosome I (Figure 5). We validated these findings by sequencing eight random clones by long-read sequencing (Supplementary Figure 4A). Although the high occurrence of additional copies for Chromosome VIII strongly suggest an adaptive strategy under galactose, populations with extra copies do not correlate with higher fitness than euploid populations (*p* = 0.56, Wilcoxon Rank-Sum test; Supplementary Figure 4B). In addition to loss or gain of entire chromosomes, we also detect large (>40 kb) CNVs in our evolved clones, including eight unique amplifications and two deletions (Table 2 and Supplementary Figure 5). Although CNVs are a common adaptive mechanism (Gorkovskiy and Verstrepen 2021), we do not find recurrent CNVs associated with galactose metabolism (Table 2 and Supplementary Figure 5). Although we found several CNVs in the right arm of Chromosome II, in only population A4 does the amplification include the *GAL1*-*GAL10*-*GAL7* gene cluster (chrII: 268984-275164). In population A4 and B9, we found duplication of genes associated with Ras/PKA signaling pathway; *IRA1* (chrII: 534751-544029) and *RAS2* (chrXIV: 440898-441866), respectively. Additionally, we found amplifications of regions containing hexose transporters genes; *HXT14* (population B9), *HXT9, HTX8, HXT16* (population G4), *HXT4, HXT1*, and *HXT5* (populations with aneuploidies for Chromosome VIII). These data demonstrate recurrent amplifications of regions that codified hexose transporters, suggesting a common adaptive mechanism in our galactose evolution.

**Figure 5.**
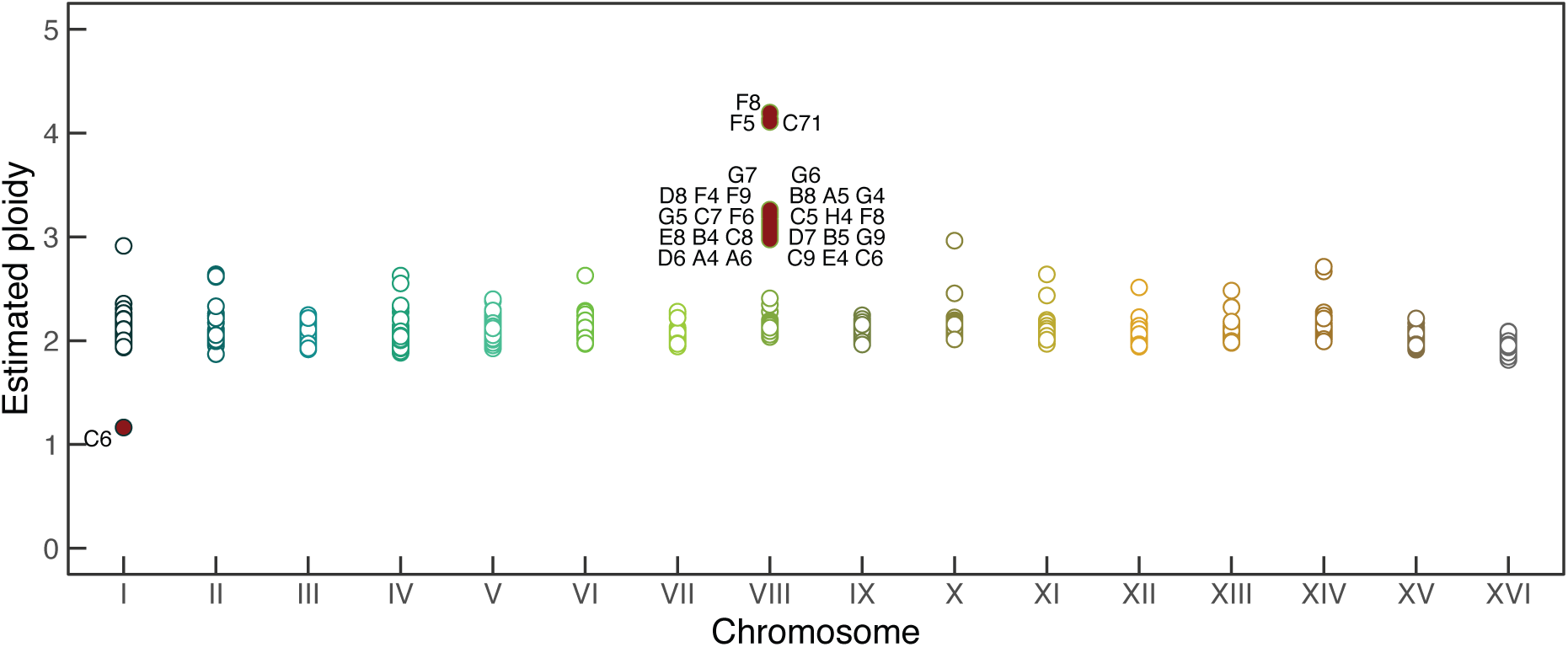
Aneuploidy in chromosome VIII appears to be a parallel evolutionary adaptation in the galactose condition. To estimate the ploidy of 96 clones, we calculated the median coverage across each chromosome compared to genome-wide coverage. Baseline ploidy is 2N. Aneuploidies are shown as filled red circles and labeled by name population. Empty circles indicate euploidy. Notably, duplication of Chromosome VIII occurred in 28 populations out of 46 populations.

**Table 2.** Structural variants in galactose-evolved populations

| Chr. | Start (kb) | Stop (kb) | Length (kb) | Copy number | Class | Clones |
| --- | --- | --- | --- | --- | --- | --- |
| I | 0 | 161000 | 161 | 1N | aneuploidy | **C61, C71** |
| II<sup>1,3</sup> | 226000 | 844051 | 618 | 3N | CNV | **A41, A42** |
| II | 666000 | 836000 | 170 | 3N | CNV | **B61, B62** |
| II<sup>3</sup> | 354000 | 836000 | 482 | 3N | CNV | **H71, H72** |
| II | 737000 | 844051 | 107 | 1N | CNV | **G71, G72** |
| VII | 749000 | 1092105 | 343 | 3N | CNV | **D51, D52** |
| VIII<sup>2</sup> | 0 | 844051 | 844 | 3N & 4N | aneuploidy | **G7,G6,D8,F4,F9,B8,A5,G4,G5,C7,F6,C5,H4,F8,E8,B4,C8,D7,B5,G9,D6,A4,A6,C9,E4,C6, D9** |
| X<sup>2</sup> | 502000 | 762303 | 260 | 4N | CNV | **G41,G42** |
| XI | 0 | 297000 | 297 | 3N & 4N | CNV | **H91, H92, H82** |
| XIII | 379000 | 750000 | 371 | 3N | CNV | F52 |
| XIII | 0 | 44000 | 44 | 3N | CNV | **D41, D42** |
| XIV<sup>2,3</sup> | 0 | 479001 | 479 | 3N | CNV | **B91, B92** |

Bolded clones indicate the SNP was found in all clones of the population
<sup>1</sup> Duplication of the GAL cluster. Coordinates: chrII. 268984-275164, W303 ref.
<sup>2</sup> Duplication of hexose transporters
<sup>3</sup> Duplication of components of the Ras/PKA pathway.

### Changes in copy number of extrachromosomal elements and reduction of the killer phenotype in galactose-evolved populations

In addition to evolution in the nuclear genome, we determined the extent to which evolution in rich galactose medium impacts three cytoplasmic element: the mitochondrial genome, the 2-micron plasmid, and the yeast dsRNA Killer virus. Using read depth as a proxy, we estimated the copy number of mitochondrial DNA for each clone according to Chiara et al. 2020. Compared to glucose-evolved population, where the mitochondrial DNA is maintained throughout evolution (Supplementary figure 7C), we found that only six galactose-evolved populations maintained or increased the copy number compared to the ancestor (19:1 ratio between mitochondrial and nuclear genomes). Most of the populations have a significant decrease in the copy number of the mitochondrial genome (*p*<0.0001, Wilcoxon rank-sum test), with a global mean of ∼8:1 mitochondrial to nuclear genomes (Supplementary figure 6A). The copy number in mitochondria estimation is highly correlated between clones from the same population (*r* = 0.928, *p* <0.0001, Pearson correlation; Supplementary Figure 7A).

In addition, we estimated the copy number of the extrachromosomal 2-micron plasmid and the reduction of the killer-associated phenotype. Our galactose-evolved populations show a drastic loss of 2-micron copy number. It was reduced from 133 initial copies to 57 copies on average per every copy of the nuclear genome. Only three populations showed a higher 2-micron copy number than the ancestor (Supplementary figure 6B). We did not observe differences in reduction of 2-micron copy number between galactose and glucose-evolved populations (*p*=0.43, Wilcoxon rank-sum test; Supplementary Figure 7D). Like mitochondrial copy number, our estimates of 2-micron copy number are highly correlated between clones from the same population (*r* = 0.66, *p* <0.0001, Pearson correlation; Supplementary Figure 7B). We did not find correlation between reduction of mitochondrial DNA and 2-micron (*r*=-0.07, *p*=0.48, Pearson correlation). As a proxy for the loss of the killer toxin, we quantified killing ability in our populations (Buskirk et al. 2020). Similar with glucose evolution, we find that the majority of the populations (42/48) lost totally or partially the killer-associated phenotype (Supplementary Figure 6C, Supplementary Figure 8). Consistent with previous glucose evolution, reduction of the 2-micron and killing ability is also a common mechanism in our galactose evolution adaptation. In contract, mitochondrial DNA has recurrent losses only in galactose evolution.

## DISCUSSION

The genetic switch from glucose to galactose utilization in *S. cerevisiae* has been extensively studied (Acar et al. 2005; Ronen and Botstein 2006; Brink et al. 2009; Escalante-Chong et al. 2015). However, the strategies for long-term adaptation to galactose as a sole carbon source are not well understood. To determine how galactose affects adaptation in budding yeast, we evolved 48 diploid populations in rich galactose medium under identical conditions to our previous glucose evolution experiment (Marad et al. 2018). We show that fitness gains over the first 2,000 generations were greater in the galactose medium compared to glucose. Because growth is slower in galactose than in glucose, some of the fitness gains could be attributed to general diminishing-returns epistasis, where beneficial mutations exert a larger fitness benefit in populations that are further from the fitness optimum (Kryazhimskiy et al. 2014). Indeed, we do find that beneficial mutations in the galactose-evolved populations are slightly beneficial in glucose medium. Similarly, glucose-evolved mutations are strongly beneficial in glucose and, but less so, beneficial in galactose, suggesting that environment-specific selective pressures drive the fixation of specific mutations in each environment.

To identify the targets of selection during long-term growth on galactose, we sequenced two clones from each population after 4,000 generations. Given the previous natural variation found in the *GAL* pathway, it is reasonable to expect multiple mutations in genes involved in this pathway. However, we do not observe more mutations than expected by chance in the *GAL* pathway. Reconstruction of evolved mutations found in the *GAL* cluster do not show fitness advantage in galactose (Supplementary Figure 3B). We identify candidate targets of selection outside of the canonical galactose pathways, with *SNT1, SEC23, PEP5*, and *IRA1* as the most significant hits. *IRA1* and *SNT1* are common targets across evolution experiments (Lang et al. 2013; Fisher et al. 2018; Li et al. 2018). In contrast, *SEC23* and *PEP5* have not been previously identified as common targets of selection in glucose. These four genes show enrichment of missense and nonsense mutations providing further support that they are under selection in galactose. In diploid evolution experiments, beneficial mutations are partially dominant or overdominant (Aggeli et al. 2022). Reconstruction of evolved alleles shows that *ira1-*S700P and *sec23-*R592I are partially dominant. Consistent with being partially dominant, we find that the allele *ira1-*S700P fixed as a homozygote. It is not surprising that *sec23-*R592I did not undergo loss of heterozygosity since the majority of mutations in the evolved clones are heterozygous (1,444 of 1,529 mutations). The target *IRA1*, is involved in the negative regulation of the Ras/PKA signaling pathway is one of the best-known glucose-triggered signaling cascades in *S. cerevisiae*. It is essential for cell cycle progression and life span in response to nutrient availability (Tamanoi 2011; Broach 2012). Genes in this pathway (e.g., *RAS1, RAS2, IRA1, IRA2, GPB2*) are frequent targets of selection in different adaptive laboratory evolution experiments, including in rich-glucose culture (Lang et al. 2013; Fisher et al. 2018; Johnson et al. 2021) and nutrient-limited medium (Kao and Sherlock 2008; Wenger et al. 2011; Koschwanez et al. 2013; Hong and Gresham 2014; Venkataram et al. 2016).

In addition to *IRA1*, we also find mutations in other signaling genes including four independent mutations in *IRA2* (E4, B7, B4, A9 populations), one mutation in *GPB1* (F8 population) and one mutation in *PDE2* (G8 population). All of these mutations are nonsynonymous. A previous evolution experiment in galactose batch culture identified mutations in genes involved in the Ras/PKA signaling pathway, including *RAS2* and *CYR1* (Hong et al. 2011b; Hong and Nielsen 2013). Though these genes and pathways are common targets across evolution experiments, the specific mutations are condition specific. We find that the galactose-evolved mutation, *ira1-*S700P, has a greater fitness benefit in galactose compared in glucose. Mutations identified in experimental evolution are often pleiotropic, affecting fitness (either positively or negatively) in other environments (Ostrowski et al. 2005; Jerison et al. 2020; Bakerlee et al. 2021). These results suggest that though the Ras/PKA pathway is a general hub for adaptive mutations in all laboratory evolution experiments, adjusting cell physiology match the environment.

Outside of the Ras/PKA pathway, the secretory pathway is another common target of selection in galactose. Apart from recurrent mutations in *SEC23*, we also see independent moderate-impact mutations in six more genes that encode secretory proteins. None of these genes are observed in our glucose-evolved populations (Marad et al. 2018). However, despite *SNT1* being a common target of selection, two of our reconstructed *SNT1* mutations did not confer a selective advantage on their own, or they were deleterious (Figure 4). It is possible the presence of epistasis interactions between *SNT1* mutations and the genetic background. We noticed that the *snt1*-A1009S and *snt1-*V237I mutations arose in populations with mutations in *ira1* and *ira2*, respectively. Taken together this data with previous evolution experiments, secretion, like nutrient sensing and signaling, is a common target of selection in galactose medium.

Previous studies show that aneuploidies are a frequent adaptive mechanism (Hong and Gresham 2014; Sunshine et al. 2015). Aneuploidies and copy number variants (CNVs) could provide a benefit by increasing the copy number of specific genes whose expression level is limiting in a particular environment (Sunshine et al. 2015). For example, the amplification of transporter and specific permeases are a recurrent adaptive mechanism to selection in medium limited for glucose (Gresham et al. 2008; Kao and Sherlock 2008), nitrogen (Hong and Gresham 2014), sulfur (Hong and Gresham 2014; Sunshine et al. 2015), or raffinose (Scott et al. 2017). An essential step in utilizing galactose is its transport achieved by Gal2p, a closely related hexose transporter (Boles and Hollenberg 1997). Notably, we did not detect any duplications of the transporter *GAL2* located on Chromosome XII. Of the four CNVs on Chromosome II, only the largest one encompasses the *GAL1*-*GAL10*-*GAL7* gene cluster (Supplementary Figure 5). We do, however, find that 60% of our populations have fixed aneuploidies on Chromosome VIII. We suggest that a strong adaptive advantage in our populations can be explained by the amplification of hexose transporters on Chromosome VIII: *HXT4, HXT1, HXT5*. Under galactose conditions, selection for spontaneous duplication of Chromosome VIII may favor nutrient transporters, leading to more efficient nutrient uptake into central metabolism (Torres et al. 2007). We only detected one point mutations in any of the *HXT* genes (a single missense mutation in *HXT3*).

Most of the observed adaptive processes have focused on the nuclear genome. Little is known about the evolution of extrachromosomal elements in different environments. Changes in copy number can give us a first light of whether these elements are under selection or not. Recently, Johnson *et al*. 2020 showed the recurrent loss of 2-micron plasmids and reduction of the killer phenotype of the Killer virus under rich glucose and synthetic complete media. Our galactose-evolved populations show a similar pattern in 2-micron plasmids and Killer-associated phenotypes (Supplementary Figure 6).

In contrast with glucose evolution, we observe multiple losses of mitochondrial DNA (Supplementary Figure 6). Under galactose metabolism, *S. cerevisiae* does not exhibit a Kluyver effect (inability to grow on and ferment sugars under anaerobic conditions). Therefore, it can consume galactose under respiration and fermentation (Brink et al. 2009). The existence of this respiration-dependent assimilation of sugars is an essential link between mitochondria and sugar utilization (Quarterman et al. 2016). The reduction of genome copy number may indicate that the evolved populations are increasing fermentation and decreasing respiration.

Overall, we find that long-term adaptation to growth on galactose is driven mainly by mutations on nutrient sensing and signaling. Specifically, we identify Ras/PKA signaling as a major target of selection, as it is in nearly all environments. We find that glucose and galactose-evolved alleles are more beneficial in the condition in which they arose. Unlike in glucose-evolved populations, we find recurrent aneuploidy for Chromosome VIII suggestion a potential role in galactose adaptation by increasing the copy number of *HXT* genes. We observe a reduction in 2-micron copy number, mitochondrial copy number, and killing ability across our evolved populations.

## METHODS

### Strains construction

All strains used in this study are derived from the W303 background (*ade2-1, CAN1, his3-11, leu2-3,112, trp1-1, URA3, bar1Δ::ADE2, hmlαΔ::LEU2, GPA1::NatMX*, ura3Δ::pFUS1-yEVenus). All the derivative strains are identified by their yGIL prefix and the number in the Lang Lab yeast collection. Mutant strains were constructed using CRISPR-Cas9 as described previously (Fisher et al. 2019). Repair oligonucleotide templates were generated by amplifying 500bp gBlocks (IDT) that contained the point mutation and a synonymous PAM site substitution. We co-transformed a high-copy plasmid that contains the Cas9 gene and the guide RNA expression cassette (pML104; Addgene #67638) (Laughery et al. 2015) and the linear repair template into yGIL432 (*MAT*a) and yGIL646 (*MAT*α) strains. We designed the following sgRNA; *snt1-*A168S (5’
s-TGGAGCGTTT GATAATGCCG AGG-3′), *snt1*-R1009K (5′-ATGGCTCTAT AAGACCATTT GGG-3′), *sec23-*R592I (5′-TTGGGATCTT CTTAAATAAT AGG-3′), and *ira1-*S700P (5′-GAAGAGATTC TACAACTTGT TGG-3′). After editing, we removed the plasmid on media containing 5-FOA. Diploid mutants were generated by crossing each mutant strain with the opposite mating type mutant strain or wild-type strain to produce homozygotes and heterozygotes, respectively. Crosses were sporulated to confirm diploid strains. All plasmids and strain construction were confirmed by Sanger sequencing (Genscript).

### Evolution experiment

Long-term evolution was performed as described previously (Marad et al. 2018). Briefly, the ancestral diploid strain, yGIL672, was grown to saturation in YPD (yeast extract, peptone, dextrose) medium. A dilution of 1:2^10^ was used to seed 48 replicate populations in a single round bottom 96-well plate. The cultures were propagated for 4,000 generations in YPGal medium (yeast extract, peptone, 2% galactose) containing ampicillin (100 mg/mL) and tetracycline (25 mg/mL) to prevent bacterial contamination. The cultures were incubated at 30°C in an unshaken 96-well plate. Every 24 h, the populations were diluted 1:1,024 by serial dilution 1:32 (4 μl into 125 μl) × 1:32 (4 μl into 125 μl) into a new YPGal 96-well plate. This regime corresponds to 10 generations of growth per day at an effective population size of ∼10^5^. The long-term experimental evolution was performed using the Biomek Liquid Handler. Approximately every 50 generations, populations were cryo-archived in 15% glycerol at -80°C.

### Competitive fitness assays

To measure the effect of evolved mutations we used flow-cytometry-based competitive fitness assay previously described in Buskirk *et al*. 2017. We used for the competition the reference strains; yGIL519 (*MAT*a), yGIL699 (*MAT*α), and yGIL702 (*MAT*a/α). These strains have the ancestral background with constitutive ymCitrine integrated at the *URA3* locus. For all the evolved alleles, we performed the competition in YPD and YPGal under identical conditions. Diploid and haploid strains were competed in independent 96-well plates in an identical fashion to the evolution experiment. The reference and the experimental strains were mixed 1:1 at Generation 0 using a Biomek Liquid handler. The assays were performed for 50 generations and sampled every 10 generations. To measure density using flow-cytometry (BD FACSCanto II), we transferred 4 μL of each sample into 60μL of PBS and stored at 4 °C for 1 day. Data were analyzed in FlowJo 10.3. The selective coefficient was calculated as the slope of the change in the natural log ratio between query and reference strains. For each fitness measurement, we performed between 4-6 independent biological replicates and one technical replicate.

### Whole-Genome Sequencing

Each population, from generation 4,000, was struck to singles colonies on YPGal and two clones were isolated for sequencing. Clones were grown to saturation in YPGal and total genomic DNA was isolated for each sample using phenol-chloroform extraction and ethanol precipitation. We used the Nextera sequencing library preparation kit as described previously (Buskirk et al. 2017). Sequencing was performed on an Illumina HiSeq 2500 sequencer with 150-nucleotide paired-end reads by the Sequencing Core Facility within the Lewis-Sigler Institute for Integrative Genomics at Princeton University.

The Oxford Nanopore MinION Genomic DNA Sequencing Kit (SQK-LSK109) was used to prepare the DNA libraries of 8 evolved clones. Briefly, we performed DNA extraction using a QIAGEN 100/G. One microgram of DNA per sample was diluted in 48μL of water. End-repair was performed using NEBNext FFPE Repair Mix and the product was purified using 60 μl of Agencourt AMPure XP beads. End-prepped gDNA was quantified using the Qubit High Sensitivity assay. dA-tailing was performed using NEBNext dA-tailing module. A ligation reaction was then performed by adding 2.5μL of the Native Barcode (EXP-PBC001 and EXP-PBC096) and 25μL of Blunt/TA Ligase Master Mix. The adapter-ligated DNA was purified using Agencourt AMPure XP beads and washed with long fragment buffer. DNA was quantified using a Qubit and then loaded on to Minion R9.4.1 (product code FLO-MIN106D R9) flow cells.

### Sequencing Analysis pipeline

Raw Illumina sequencing data were concatenated and demultiplexed using barcode_splitter 0.18.6 from L. Parsons (Princeton University). Adapter sequences were trimmed using trimmomatic/0.36 (Bolger et al. 2014) using PE -phred33 parameter. Each sample was aligned to the complete and annotated W303 genome (Matheson et al. 2017) using BWA-MEM, v.0.7.15 (Li and Durbin 2009). Each clone was sequenced to an average depth of approximately 50X coverage (Supplementary Figure 1). Common variants were called using FreeBayes/1.1.0 (Garrison and Marth 2012), using default parameters. Variants common were filtered using the VCFtools/0.1.15 vcf-isec function (v.0.1.12b). Individual VCF files were annotated using SnpEff/5.0 using -formatEff flag (Cingolani et al. 2012). Manually we did a more punctual filtering of common variants by viewing BAM files using Integrated Genome Viewer (Broad Institute). We removed the variant calls found in low complexity regions (TY elements, centromeres, and telomeric regions) and with less than 5X coverage. Only nuclear mutations were analyzed. We used R package *idiogramFISH* (Roa and PC Telles. 2021) to plot the distribution of our evolved mutations. Zygosity was determined by establishing a cut-off of mutation frequency values above 0.9 and *p-*values < 0.001. If both parameters were satisfied, we called those mutations homozygous. Copy Number Variants (CNVs) were identified using ControlFREEC version 11.5 (Boeva et al. 2012). All the .txt output were merged into one data frame to remove all the common calls. We applied the following criteria to filtered CNVs: the variant is calling more than 10 times, had length less than 5 kb, and variants that were located close to telomeres and centromeres (5kb of distance) regions. CNVs and aneuploidies were corroborated by visual inspection of chromosome coverage plots created in R. Briefly, we used samtools-depth to calculate per-site depth from the sorted-bam files. We divided the median chromosome coverage by the median genome-wide coverage using non-overlapping 1000 bp window size and 500 nt step size. The total coverage of the genome was normalized to 2 (diploid population). The same analysis was used to estimate copy number of the 2-micron plasmid. Copy number of mitochondrial DNA was estimated using the same sliding window approach but only from the *ATP6, COX2*, and *COX3* regions (Chiara et al. 2020) to avoid overcounting the highly-repetitive sequences in the mitochondrial genome. For Nanopore sequencing, we performed basecalling and demultiplex barcode using guppy/3.1.5. The eight clones were aligned to the S288C reference using the long-read mapper ngmlr/0.2.7 (Sedlazeck et al. 2018). We used Sniffles (Sedlazeck et al. 2018) to identify structural variants. To visualize chromosome coverage, we used the same methodology described for Illumina data. For glucose-evolved populations, we performed the same analysis as galactose. However, the whole-genome sequencing was performed at Generation 2,000.

### Identification of common targets

The *p-*values of coding genes hit by a mutation just by chance were calculated using the Poisson distribution (Fisher et al. 2018). We determined the probability that chance alone explains the observed number of mutations across all 5,800 genes by assuming a random Poisson distribution of the 1,529 mutations weighted by the length of the gene across the 8,453525 bp genome-wide CDS. Common targets of selection were defined as genes with three or more CDS mutations and a corresponding probability of less than 0.5% and 0.1%. To determine if the targets of selection are unique to galactose, we compare the *p-*values with prior glucose evolution experiments (Marad et al. 2018). To determine targets of selection for each condition, we calculated the ratio between probabilities (Supplementary Data).

### Halo assay

Killer phenotype was performed using the protocol described previously (Buskirk et al. 2020). Assay was performed using YPD agar that had been buffered to pH 4.5 (citrate-phosphate buffer), dyed with methylene blue (0.003%), and poured into a 1-well rectangular cell culture plate. Killing ability was assayed against a hypersensitive tester strain (yGIL1097). The hypersensitive tester was grown to saturation, diluted 1:10, and spread (150 ml) evenly on the buffered agar. Glucose and galactose-evolved populations were grown to saturation, concentrated five-fold, and spotted (2 μL) using liquid handler (Biomek FX). Plates were incubated at room temperature for 3 days before assessment. Killer phenotype was scored according to the scale in shown in Buskirk et al. 2020.

### Statistical Analyses

All statistical analyses reported were performed using tools in the R Stats package in R v.4.0.2. All plots were produced in R using the ggplot2 package (Wickham 2016).

## DATA AVAILABILITY

The short-read sequencing data reported in this study have been deposited to the NCBI BioProject database, accession number PRJNA835840. The previously published dataset from Marad et al. 2018 is available under NCBI BioProject PRJNA418180.

## ACKNOWLEDGEMENTS AND FUNDING

We thank members of the Lang Lab for comments on the manuscript. This study was supported by a grant from the National Institutes of Health (R01GM127420). Portions of this research were conducted on Lehigh University’s Research Computing infrastructure partially supported by the National Science Foundation (Award 2019035).

## AUTHOR CONTRIBUTIONS

Conceptualization: GIL

Formal Analysis: AAM, GIL

Investigation: AAM, AC, DM, SWB

Writing – Original Draft Preparation: AAM, GIL

Writing – Review & Editing: AAM, GIL

Visualization: AAM, GIL

Supervision: GIL Funding Acquisition: GIL

---

**Supplementary Figure 1.**
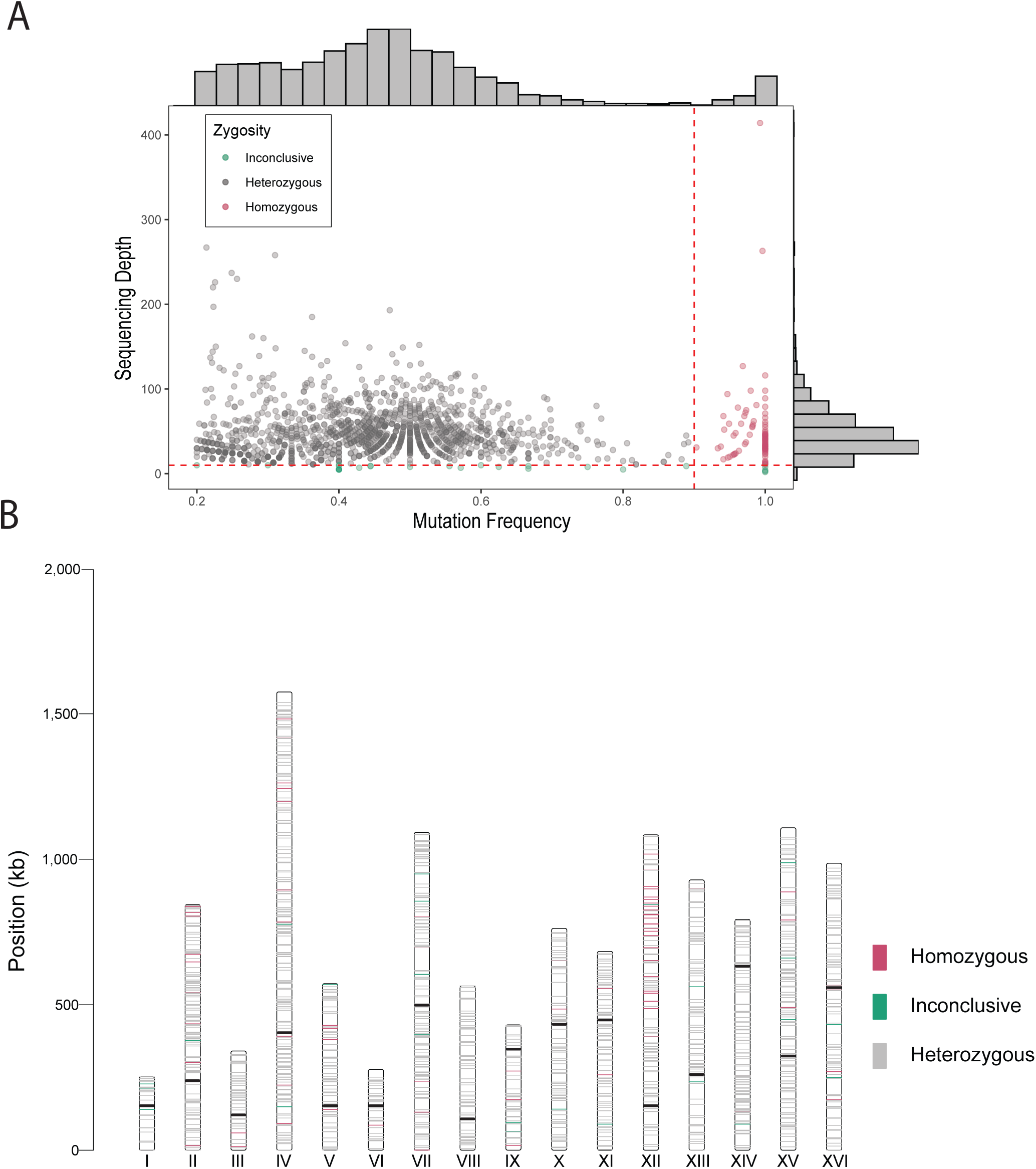
Zygosity of the evolved mutations. (A) Criteria to classify heterozygous and homozygous mutations. On the x-axis is the allele frequency and on the y-axis is the read depth for each variant called (mean=49X). The red dotted line indicates the cut-off to call homozygous mutations with *p*-values < 0.001. Only 6% of the evolved mutations are homozygotes. We call inconclusive variant if the sequencing depth is below 0.9. (B) Distribution of evolved mutations colored by zygosity. There is an enrichment of homozygous mutations in chromosome XII. Centromers are represented with a black horizontal line.

**Supplementary Figure 2.**
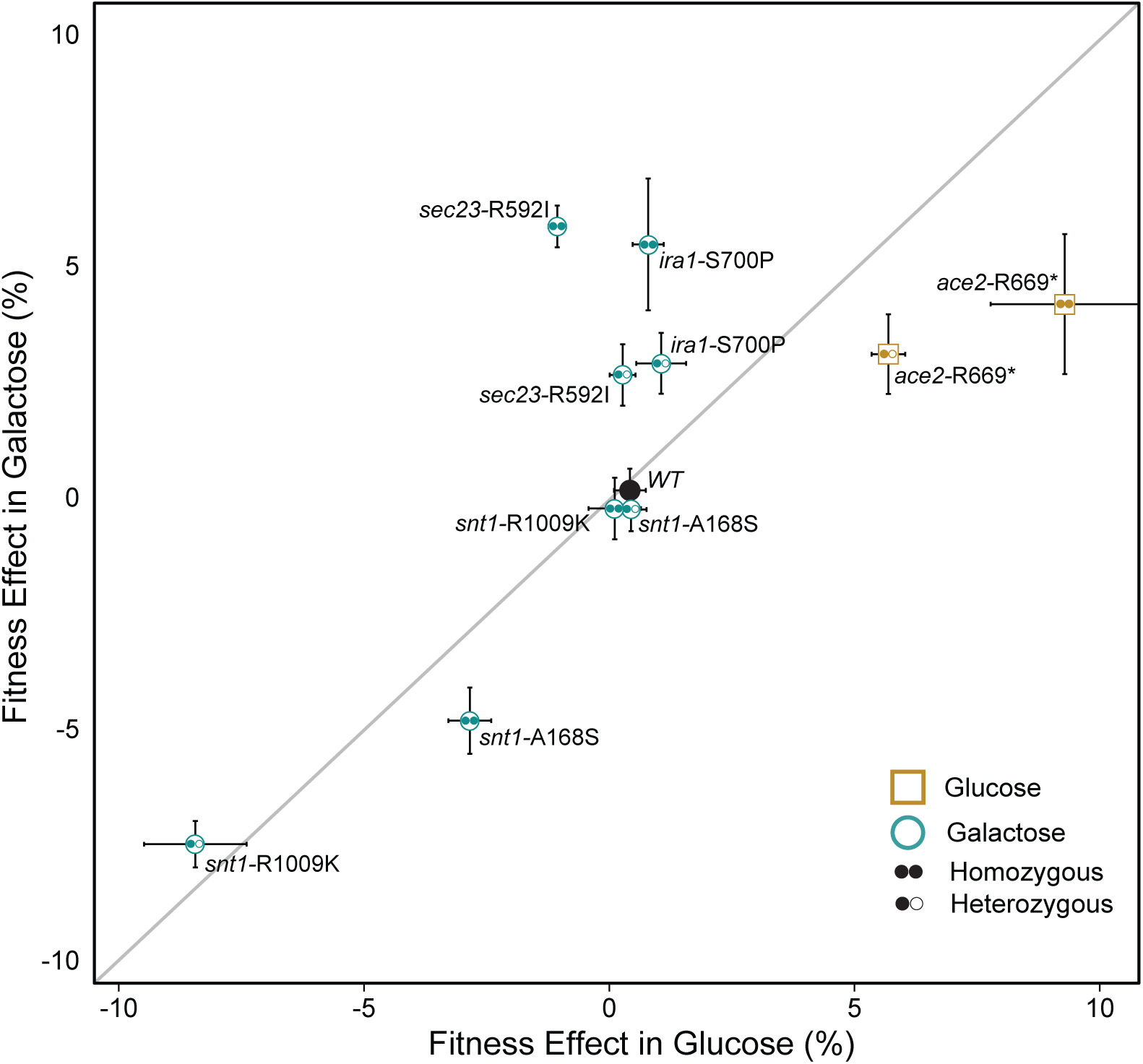
Fitness of evolved mutations correlate with the condition in which the specific mutation arose. Plotted is the correaltion of the average fitness effects of the reconstructed mutations *snt1* (n=6), *sec23* (n=6), *ira1* (n=4), and *ace2* (n=4) in heterozygous diploid and homozygous diploids in galactose and glucose condition. Mutations that arose in galactose evolution are indicated by cyan circles and mutation that arose in glucose evolution, *ace2*, is indicated by gold square. Pearson correlation coefficient *r* = 0.69, *p* = 0.0122. Error bars represent s.e.m.

**Supplementary Figure 3.**
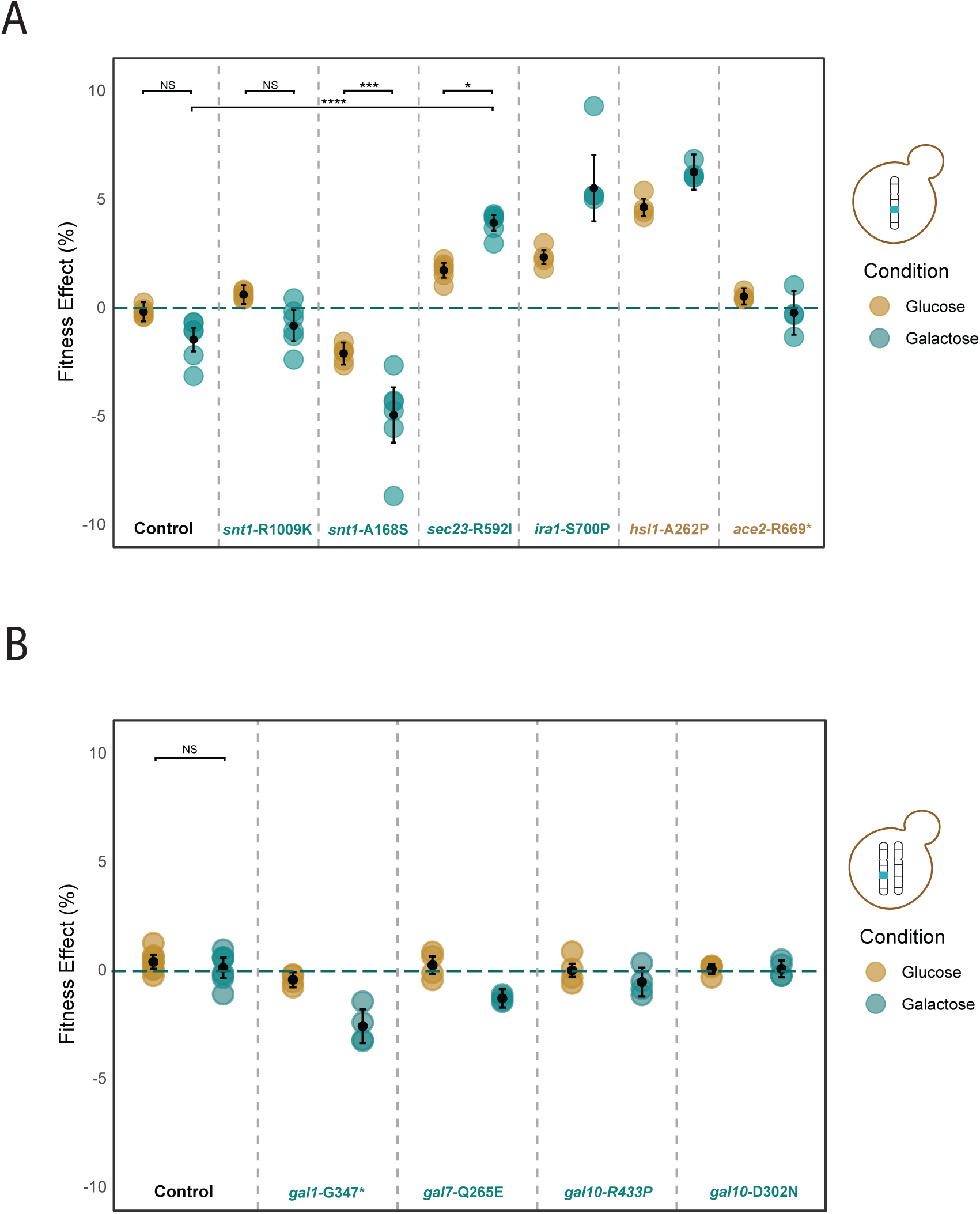
Fitness effect of Galactose-evolved alleles. (A)Average fitness effects of the reconstructed mutations in *snt1* (n=6), *sec23* (n=6), and *ira1* (n=4), *hsl1* (n=4), and *ace2* (n=4) in haploid background. The *sec23* allele has a significant fitness gain in galactose compare to glucose. Asterisk (****) indicates *p* < 10^−8^, (***) *p* < 10^−4^, (*) *p* < 10^−2^, and NS: not significant; one-way ANOVA, tukey’s HSD test. The *ira1* allele follows the same effect as *sec23* allele. (B) Average fitness effect of evolved mutations in *gal1* (n=4), *gal7* (n=4), *gal10* (n=4) in in heterozygous allele. No fitness advantage from mutations in galactose genes. We estimate *p-*value only for alleles with 6 replicates. Error bars are the s.e.m.

**Supplementary Figure 4.**
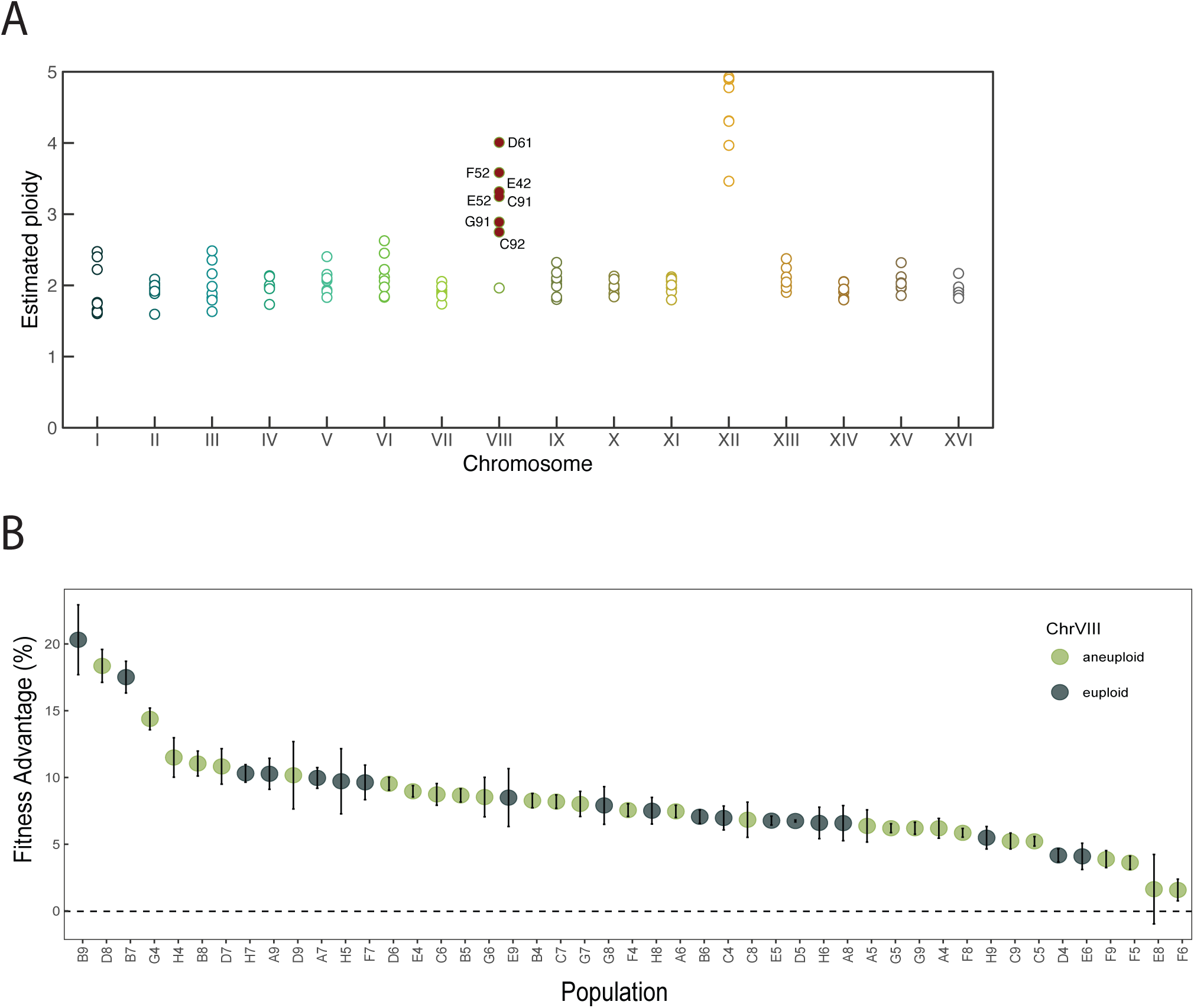
Aneuploidy in Chromosome VIII does not show correlation with populations fitness gain. (A) Ploidy estimation of 8 independent clones from Nanopore sequencing. A large peak of coverage is observed on chromosome XII, which represents the tandemly repeated regions of the rDNA loci. Aneuploidies are shown as filled red circles and labeled by name population. Empty circles indicate euploidy. (B) Fitness effect of 48 populations after 2,000 generations. Populations with 3N or 4N copies on Chromosome VIII are represented in light green circles. Dark green circles indicate euploidy for Chromosome VIII. Populations with trisomy for Chromosome VIII did not show higher fitness advantage (*p*-value = 0.56, Wilcoxon Rank-Sum test).

**Supplementary Figure 5.**
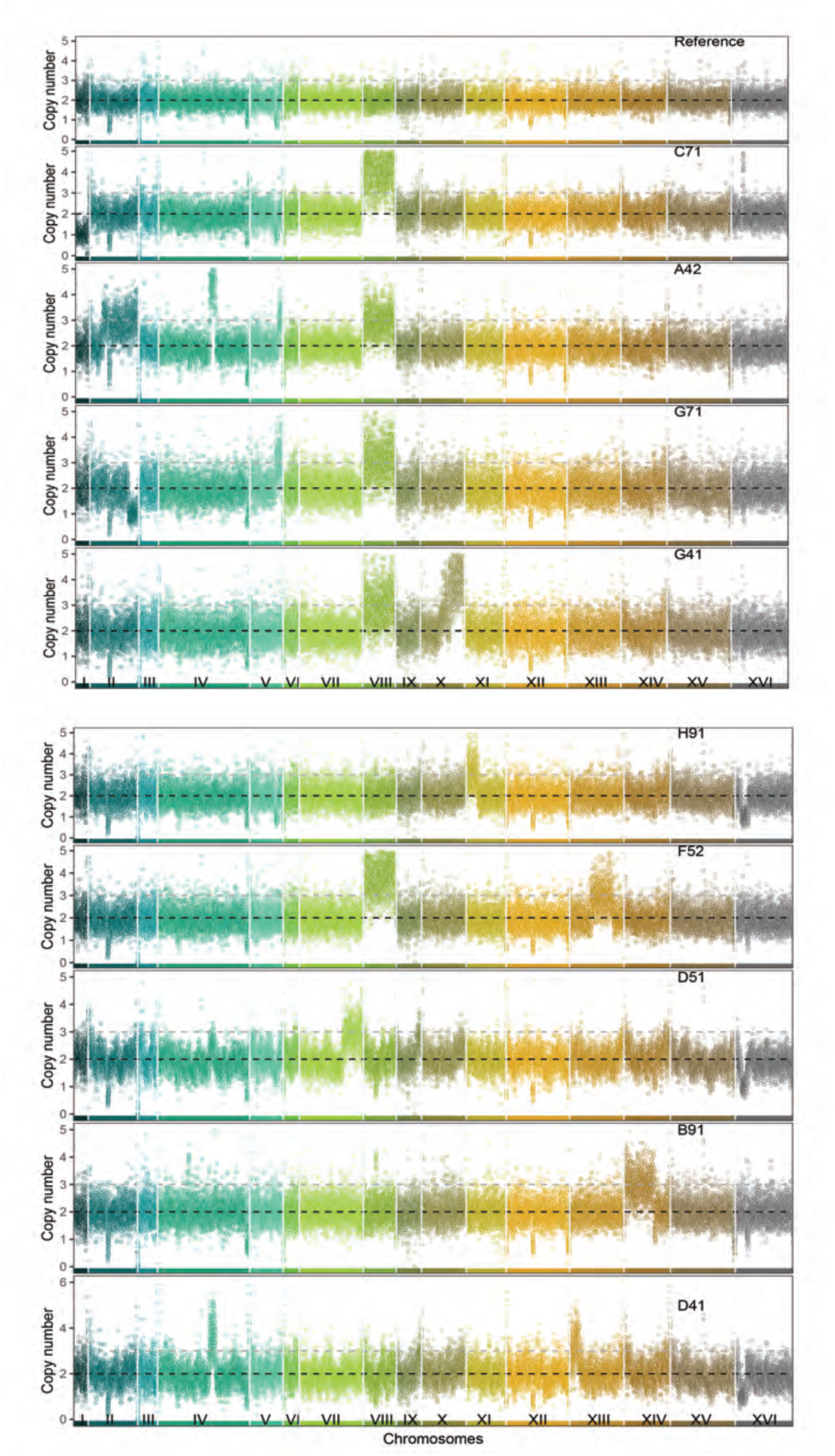
Detection of Structural Variants of nine evolved clones. Coverage across each position on the genome using a sliding window approach; window size of 1000 bp and window step of 500 bp. Each color represents different Chromosomes. The baseline ploidy was assumed to be 2N.

**Supplementary Figure 6.**
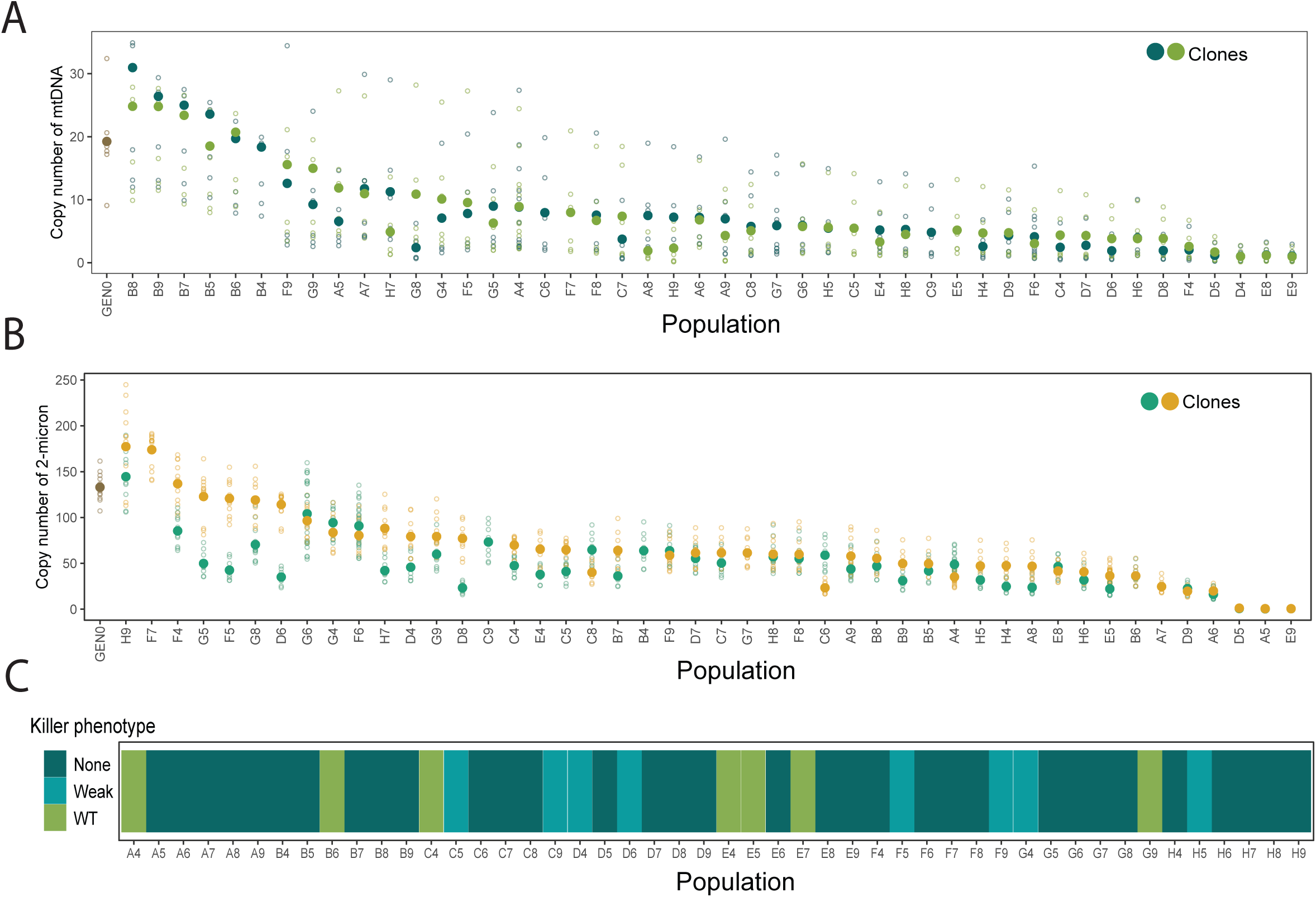
Recurrent loss of extrachromosomal elements and killing ability in galac-tose-evolved populations after 4,000 generations. Estimated copy number of (A) mitochondrial DNA and (B) 2-micron plasmid per every copy of the nuclear genome per clone. Each population is represented by the two sequenced clones labed with different colors. Open circles indicate copy number stimation every 1000 bp window size and 500 nt step size. Solid circles indicate the mean copy number. (C) Reduction of killing ability in yeast that harbor killer viruses. The killing ability was determined by Halo assay (WT=large clearing=, weak=small clearing, and none=no clearing).

**Supplementary Figure 7.**
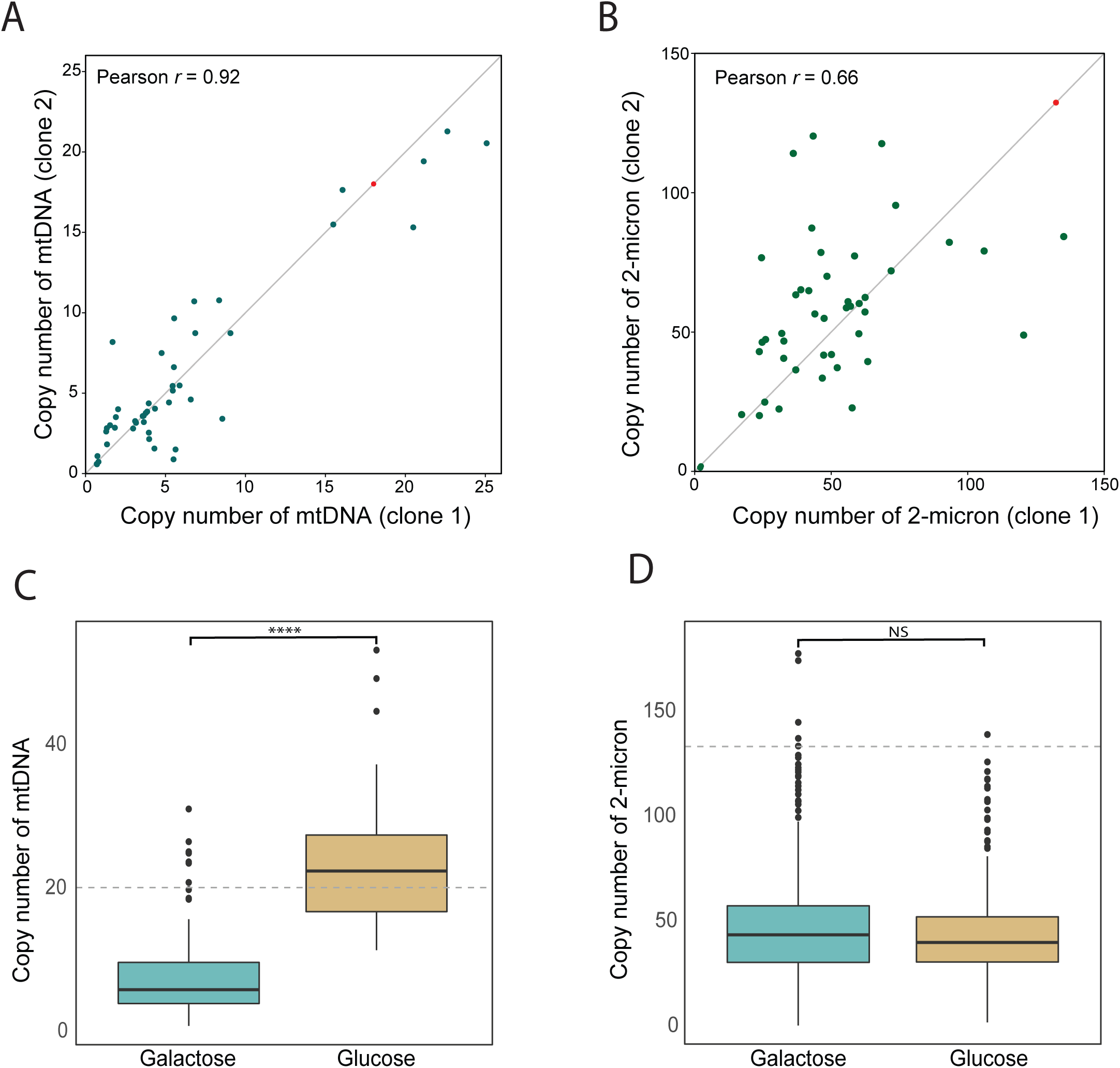
Copy number estimation of extrachromosomal elements between two clones per population and between two conditions. (A) Average copy number correlation of (A) mito-chondrial DNA and (B) 2-micron plasmid between two clones. Pearson correlation for each clone pair is shown at the top left of each panel. The ancestor is represented by a red circle. Box plot reflects the mean copy number estimation of (C) mitochondrial DNA and (D) 2-micron plasmid of galactose and glucose-evolved populations. The gray dotted line indicates the initial copy number estimation from the ancestor. (C) Galactose-evolved populations have significant losses of mitochondrial DNA compared to glucose-evolved popualtions (*p* < 0.0001, Wilcoxon rank-sum test). (D) No significant difference in losses of 2-micron between both sugars (*p* =0.43, Wilcoxon rank-sum test).

**Supplementary Figure 8.**
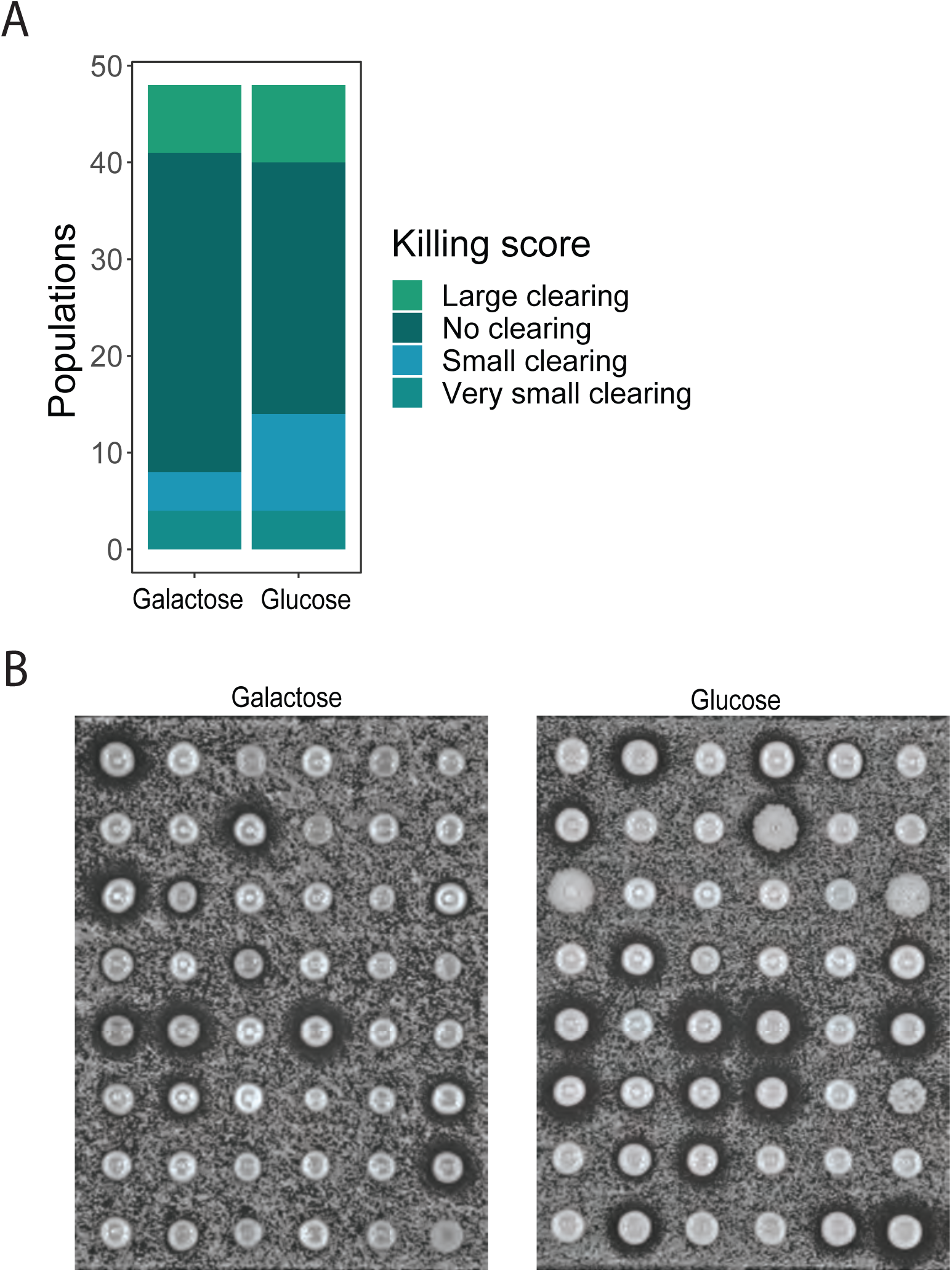
Killer-associated phenotype in galactose and glucose-evolved population. (A) Bar plot of killing score of 48 galactose and 48 glucose-evolved populations. (B) Halo assay of both evolved population.

## REFERENCE

Acar M, Becskei A, Oudenaarden A van (2005) Enhancement of cellular memory by reducing stochastic transitions. Nature 435:228–232. https://doi.org/10.1038/nature03524

Acosta PB, Gross KC (1995) Hidden sources of galactose in the environment. Eur J Pediatr 154:S87–S92. https://doi.org/10.1007/bf02143811

Aggeli D, Marad DA, Liu X, et al (2022) Overdominant and partially dominant mutations drive clonal adaptation in diploid Saccharomyces cerevisiae. Genetics. https://doi.org/10.1093/genetics/iyac061

Bakerlee CW, Phillips AM, Ba ANN, Desai MM (2021) Dynamics and variability in the pleiotropic effects of adaptation in laboratory budding yeast populations. Elife 10:e70918. https://doi.org/10.7554/elife.70918

Boeva V, Popova T, Bleakley K, et al (2012) Control-FREEC: a tool for assessing copy number and allelic content using next-generation sequencing data. Bioinformatics 28:423–425. https://doi.org/10.1093/bioinformatics/btr670

Boles E, Hollenberg CP (1997) The molecular genetics of hexose transport in yeasts. Fems Microbiol Rev 21:85–111. https://doi.org/10.1111/j.1574-6976.1997.tb00346.x

Bolger AM, Lohse M, Usadel B (2014) Trimmomatic: a flexible trimmer for Illumina sequence data. Bioinformatics 30:2114–2120. https://doi.org/10.1093/bioinformatics/btu170

Boocock J, Sadhu MJ, Durvasula A, et al (2021) Ancient balancing selection maintains incompatible versions of the galactose pathway in yeast. Science 371:415–419. https://doi.org/10.1126/science.aba0542

Brink J van den, Akeroyd M, Hoeven R van der, et al (2009) Energetic limits to metabolic flexibility: responses of Saccharomyces cerevisiae to glucose–galactose transitions. Microbiology+ 155:1340–1350. https://doi.org/10.1099/mic.0.025775-0

Broach JR (2012) Nutritional Control of Growth and Development in Yeast. Genetics 192:73–105. https://doi.org/10.1534/genetics.111.135731

Buskirk SW, Peace RE, Lang GI (2017) Hitchhiking and epistasis give rise to cohort dynamics in adapting populations. Proc National Acad Sci 114:8330–8335. https://doi.org/10.1073/pnas.1702314114

Buskirk SW, Rokes AB, Lang GI (2020) Adaptive evolution of nontransitive fitness in yeast. Elife 9:e62238. https://doi.org/10.7554/elife.62238

Chiara MD, Friedrich A, Barré B, et al (2020) Discordant evolution of mitochondrial and nuclear yeast genomes at population level. Bmc Biol 18:49. https://doi.org/10.1186/s12915-020-00786-4

Cingolani P, Platts A, Wang LL, et al (2012) A program for annotating and predicting the effects of single nucleotide polymorphisms, SnpEff. Fly 6:80–92. https://doi.org/10.4161/fly.19695

Conrad M, Schothorst J, Kankipati HN, et al (2014) Nutrient sensing and signaling in the yeast Saccharomyces cerevisiae. Fems Microbiol Rev 38:254–299. https://doi.org/10.1111/1574-6976.12065

Duan S-F, Han P-J, Wang Q-M, et al (2018) The origin and adaptive evolution of domesticated populations of yeast from Far East Asia. Nat Commun 9:2690. https://doi.org/10.1038/s41467-018-05106-7

Escalante-Chong R, Savir Y, Carroll SM, et al (2015) Galactose metabolic genes in yeast respond to a ratio of galactose and glucose. Proc National Acad Sci 112:1636–1641. https://doi.org/10.1073/pnas.1418058112

Fendt S-M, Sauer U (2010) Transcriptional regulation of respiration in yeast metabolizing differently repressive carbon substrates. Bmc Syst Biol 4:12. https://doi.org/10.1186/1752-0509-4-12

Fisher KJ, Buskirk SW, Vignogna RC, et al (2018) Adaptive genome duplication affects patterns of molecular evolution in Saccharomyces cerevisiae. Plos Genet 14:e1007396. https://doi.org/10.1371/journal.pgen.1007396

Fisher KJ, Kryazhimskiy S, Lang GI (2019) Detecting genetic interactions using parallel evolution in experimental populations. Philosophical Transactions Royal Soc B 374:20180237. https://doi.org/10.1098/rstb.2018.0237

Garrison E, Marth G (2012) Haplotype-based variant detection from short-read sequencing. Arxiv

Gorkovskiy A, Verstrepen KJ (2021) The Role of Structural Variation in Adaptation and Evolution of Yeast and Other Fungi. Genes-basel 12:699. https://doi.org/10.3390/genes12050699

Gresham D, Desai MM, Tucker CM, et al (2008) The Repertoire and Dynamics of Evolutionary Adaptations to Controlled Nutrient-Limited Environments in Yeast. Plos Genet 4:e1000303. https://doi.org/10.1371/journal.pgen.1000303

Harrison M-C, LaBella AL, Hittinger CT, Rokas A (2021) The evolution of the GALactose utilization pathway in budding yeasts. Trends Genet 38:97–106. https://doi.org/10.1016/j.tig.2021.08.013

Hittinger CT, Carroll SB (2007) Gene duplication and the adaptive evolution of a classic genetic switch. Nature 449:677–681. https://doi.org/10.1038/nature06151

Hong J, Gresham D (2014) Molecular Specificity, Convergence and Constraint Shape Adaptive Evolution in Nutrient-Poor Environments. Plos Genet 10:e1004041. https://doi.org/10.1371/journal.pgen.1004041

Hong K-K, Nielsen J (2013) Adaptively evolved yeast mutants on galactose show trade-offs in carbon utilization on glucose. Metab Eng 16:78–86. https://doi.org/10.1016/j.ymben.2013.01.007

Hong K-K, Vongsangnak W, Vemuri GN, Nielsen J (2011a) Unravelling evolutionary strategies of yeast for improving galactose utilization through integrated systems level analysis. Proc National Acad Sci 108:12179–12184. https://doi.org/10.1073/pnas.1103219108

Hong K-K, Vongsangnak W, Vemuri GN, Nielsen J (2011b) Unravelling evolutionary strategies of yeast for improving galactose utilization through integrated systems level analysis. Proc National Acad Sci 108:12179–12184. https://doi.org/10.1073/pnas.1103219108

Jacob F, Monod J (1961) Genetic regulatory mechanisms in the synthesis of proteins. J Mol Biol 3:318–356. https://doi.org/10.1016/s0022-2836(61)80072-7

Jerison ER, Ba ANN, Desai MM, Kryazhimskiy S (2020) Chance and necessity in the pleiotropic consequences of adaptation for budding yeast. Nat Ecol Evol 4:601–611. https://doi.org/10.1038/s41559-020-1128-3

Johnson MS, Gopalakrishnan S, Goyal J, et al (2021) Phenotypic and molecular evolution across 10,000 generations in laboratory budding yeast populations. Elife 10:e63910. https://doi.org/10.7554/elife.63910

Kao KC, Sherlock G (2008) Molecular Characterization of Clonal Interference during Adaptive Evolution in Asexual Populations of Saccharomyces cerevisiae. Nat Genet 40:1499–1504. https://doi.org/10.1038/ng.280

Koschwanez JH, Foster KR, Murray AW (2013) Improved use of a public good selects for the evolution of undifferentiated multicellularity. Elife 2:e00367. https://doi.org/10.7554/elife.00367

Kryazhimskiy S, Rice DP, Jerison ER, Desai MM (2014) Global epistasis makes adaptation predictable despite sequence-level stochasticity. Science 344:1519–1522. https://doi.org/10.1126/science.1250939

Lang GI, Botstein D (2011) A Test of the Coordinated Expression Hypothesis for the Origin and Maintenance of the GAL Cluster in Yeast. Plos One 6:e25290. https://doi.org/10.1371/journal.pone.0025290

Lang GI, Rice DP, Hickman MJ, et al (2013) Pervasive genetic hitchhiking and clonal interference in forty evolving yeast populations. Nature 500:571–574. https://doi.org/10.1038/nature12344

Laughery MF, Hunter T, Brown A, et al (2015) New vectors for simple and streamlined CRISPR–Cas9 genome editing in Saccharomyces cerevisiae. Yeast 32:711–720. https://doi.org/10.1002/yea.3098

Lee K, Hong M, Jung S, et al (2011) Improved galactose fermentation of Saccharomyces cerevisiae through inverse metabolic engineering. Biotechnol Bioeng 108:621–631. https://doi.org/10.1002/bit.22988

Lee KB, Wang J, Palme J, et al (2017) Polymorphisms in the yeast galactose sensor underlie a natural continuum of nutrient-decision phenotypes. Plos Genet 13:e1006766. https://doi.org/10.1371/journal.pgen.1006766

Legras J-L, Galeote V, Bigey F, et al (2018) Adaptation of S. cerevisiae to Fermented Food Environments Reveals Remarkable Genome Plasticity and the Footprints of Domestication. Mol Biol Evol 35:1712–1727. https://doi.org/10.1093/molbev/msy066

Li H, Durbin R (2009) Fast and accurate short read alignment with Burrows–Wheeler transform. Bioinformatics 25:1754–1760. https://doi.org/10.1093/bioinformatics/btp324

Li Y, Venkataram S, Agarwala A, et al (2018) Hidden Complexity of Yeast Adaptation under Simple Evolutionary Conditions. Curr Biol 28:515-525.e6. https://doi.org/10.1016/j.cub.2018.01.009

Marad DA, Buskirk SW, Lang GI (2018) Altered access to beneficial mutations slows adaptation and biases fixed mutations in diploids. Nat Ecol Evol 2:882–889. https://doi.org/10.1038/s41559-018-0503-9

Matheson K, Parsons L, Gammie A (2017) Whole-Genome Sequence and Variant Analysis of W303, a Widely-Used Strain of Saccharomyces cerevisiae. G3 Genes Genomes Genetics 7:2219–2226. https://doi.org/10.1534/g3.117.040022

Nair A, Sarma SJ (2021) The impact of carbon and nitrogen catabolite repression in microorganisms. Microbiol Res 251:126831. https://doi.org/10.1016/j.micres.2021.126831

New AM, Cerulus B, Govers SK, et al (2014) Different Levels of Catabolite Repression Optimize Growth in Stable and Variable Environments. Plos Biol 12:e1001764. https://doi.org/10.1371/journal.pbio.1001764

Ostrowski EA, Rozen DE, Lenski RE (2005) PLEIOTROPIC EFFECTS OF BENEFICIAL MUTATIONS IN ESCHERICHIA COLI. Evolution 59:2343–2352. https://doi.org/10.1111/j.0014-3820.2005.tb00944.x

Peng W, Liu P, Xue Y, Acar M (2015) Evolution of gene network activity by tuning the strength of negative-feedback regulation. Nat Commun 6:6226. https://doi.org/10.1038/ncomms7226

Quarterman J, Skerker JM, Feng X, et al (2016) Rapid and efficient galactose fermentation by engineered Saccharomyces cerevisiae. J Biotechnol 229:13–21. https://doi.org/10.1016/j.jbiotec.2016.04.041

Ronen M, Botstein D (2006) Transcriptional response of steady-state yeast cultures to transient perturbations in carbon source. P Natl Acad Sci Usa 103:389–394. https://doi.org/10.1073/pnas.0509978103

Scott AL, Richmond PA, Dowell RD, Selmecki AM (2017) The Influence of Polyploidy on the Evolution of Yeast Grown in a Sub-Optimal Carbon Source. Mol Biol Evol 34:2690–2703. https://doi.org/10.1093/molbev/msx205

Sedlazeck FJ, Rescheneder P, Smolka M, et al (2018) Accurate detection of complex structural variations using single molecule sequencing. Nat Methods 15:461–468. https://doi.org/10.1038/s41592-018-0001-7

Sellick CA, Campbell RN, Reece RJ (2008) Chapter 3 Galactose Metabolism in Yeast—Structure and Regulation of the Leloir Pathway Enzymes and the Genes Encoding Them. Int Rev Cel Mol Bio 269:111–150. https://doi.org/10.1016/s1937-6448(08)01003-4

Slepak T, Tang M, Addo F, Lai K (2005) Intracellular galactose-1-phosphate accumulation leads to environmental stress response in yeast model. Mol Genet Metab 86:360–371. https://doi.org/10.1016/j.ymgme.2005.08.002

Sood V, Brickner JH (2017) Genetic and Epigenetic Strategies Potentiate Gal4 Activation to Enhance Fitness in Recently Diverged Yeast Species. Curr Biol 27:3591-3602.e3. https://doi.org/10.1016/j.cub.2017.10.035

Sunshine AB, Payen C, Ong GT, et al (2015) The Fitness Consequences of Aneuploidy Are Driven by Condition-Dependent Gene Effects. Plos Biol 13:e1002155. https://doi.org/10.1371/journal.pbio.1002155

Swamy KBS, Zhou N (2019) Experimental evolution: its principles and applications in developing stress-tolerant yeasts. Appl Microbiol Biot 103:2067–2077. https://doi.org/10.1007/s00253-019-09616-2

Tamanoi F (2011) Ras Signaling in Yeast. Genes Cancer 2:210–215. https://doi.org/10.1177/1947601911407322

Torres EM, Sokolsky T, Tucker CM, et al (2007) Effects of Aneuploidy on Cellular Physiology and Cell Division in Haploid Yeast. Science 317:916–924. https://doi.org/10.1126/science.1142210

Venkataram S, Dunn B, Li Y, et al (2016) Development of a Comprehensive Genotype-to-Fitness Map of Adaptation-Driving Mutations in Yeast. Cell 166:1585-1596.e22. https://doi.org/10.1016/j.cell.2016.08.002

Vignogna RC, Buskirk SW, Lang GI (2021) Exploring a local genetic interaction network using evolutionary replay experiments. Mol Biol Evol msab087.. https://doi.org/10.1093/molbev/msab087

Voordeckers K, Kominek J, Das A, et al (2015) Adaptation to High Ethanol Reveals Complex Evolutionary Pathways. Plos Genet 11:e1005635. https://doi.org/10.1371/journal.pgen.1005635

Wang J, Atolia E, Hua B, et al (2015) Natural Variation in Preparation for Nutrient Depletion Reveals a Cost–Benefit Tradeoff. Plos Biol 13:e1002041. https://doi.org/10.1371/journal.pbio.1002041

Wenger JW, Piotrowski J, Nagarajan S, et al (2011) Hunger Artists: Yeast Adapted to Carbon Limitation Show Trade-Offs under Carbon Sufficiency. Plos Genet 7:e1002202. https://doi.org/10.1371/journal.pgen.1002202

Yona AH, Manor YS, Herbst RH, et al (2012) Chromosomal duplication is a transient evolutionary solution to stress. Proc National Acad Sci 109:21010–21015. https://doi.org/10.1073/pnas.1211150109

Zaman S, Lippman SI, Zhao X, Broach JR (2008) How Saccharomyces Responds to Nutrients. Genetics 42:27–81. https://doi.org/10.1146/annurev.genet.41.110306.130206

